# Multiple-level organizational logic of the locus coeruleus-norepinephrine system in structure and function

**DOI:** 10.1101/2025.06.11.659024

**Authors:** Chang-Mei Zhang, Yong-Xin Yang, Fu-Ning Li, Hui Zhang, Le Sun, Jia Li, Chen Yang, Zhi-Feng Yue, Jie-Si Feng, Yu-Long Li, Rong-Wei Zhang, Jie He, Xu-Fei Du, Jiu-Lin Du

## Abstract

The locus coeruleus (LC) regulates various neural processes through a limited number of norepinephrine (NE)-releasing neurons with widespread axon projections. However, how the LC is organized to exert brain-wide regulatory functions remain elusive. Utilizing larval zebrafish as a whole-brain-scale model, we reveal that morphologically heterogenous but physiologically comparable LC-NE neurons assemble complementarily to implement region-specific regulation of brain-wide neural dynamics. Individual LC-NE neurons exhibit diverse axon projections with ipsilateral bias and region preference, but share comparable electrophysiological properties, sensory responses and neural function-related genes’ expression. Moreover, population LC-NE neurons display bilaterally symmetrical yet regionally different axon projections and fire synchronously, causing regionally patterned NE release. Along with the regional comparability of NE receptor expression, these enable the LC to tune brain-wide neural dynamics in an inverted-U manner with region difference. Thus, our study provides a comprehensive understanding of the organizational logic of the LC system.

## INTRODUCTION

The locus coeruleus (LC), a compact nucleus in the brainstem, is the primary source of the neuromodulator norepinephrine (NE) in the brain.^1-6^ It is comprised of a densely packed noradrenergic neurons and evolutionarily conserved across vertebrates.^7-12^ Through its brain-wide axonal projections and associated NE release, the LC regulates a large spectrum of neural functions, including sensory processing, arousal and sleep, learning and memory, attention, and physiological stress.^11,13-25^ Dysfunction of LC-NE neurons are highly associated with cognitive deficits and neurological disorders.^26-31^ A long-lasting interesting question in the field is how the compact LC organizes itself to implement such brain-wide diverse functional regulation. In particular, it remains unclarified whether and to what extent the LC system works as a homogeneous unit or in modularity.^4,13,32,33^

The LC is traditionally considered as a homogeneous unit in its structure as well as function for decades. Densely packed LC-NE neurons receive convergent inputs with dense dendritic arborizations and broadcast information via divergent axon projections.^5,10,34,35^ These neurons derive mainly from the same progenitors,^7,36^ display correlated spontaneous burst firing and similar responses to salient stimuli, show neurochemical uniformity in releasing NE throughout the brain, and regulate neural functions under either tonic or phasic firing mode.^15,35,37-48^ These lines of evidence collectively suggests a general trend toward functional homogeneity for the LC in regulating brain neural networks, though without systematic interpretation from anatomy to function.^1-3,49,50^

With the advancement of cutting-edge technologies, emerging evidence over the past decade indicates the heterogeneity of LC-NE neurons in molecular profile, anatomy, activity, and axon projection-specific regulatory functions. Distinct subpopulations of LC-NE neurons are identified by genetic assays and single-cell sequencing.^51-54^ Barcoded RNA technology and retrograde tracing suggest anatomical heterogeneity and complexity of LC-NE neurons.^10,55-59^ *In vitro* electrophysiological recordings indicate different activity patterns between LC-NE neurons innervating the prefrontal and motor cortices,^56^ or projecting to the spinal cord and those in the core of the LC.^60^ *In vivo* electrophysiological recordings in rodents showed diverse spike waveforms and sparse synchrony within population LC-NE neurons.^61-63^ Notably, a series of functional studies integrating with retrograde labeling revealed that LC-NE neurons exhibit axon projection-specific modularity in regulating specific cognitive functions and behaviors, with target brain regions including the prefrontal cortex, motor cortex, hippocampus, amygdala, cerebellum, and olfactory bulb.^11,21,44,58,59,61,64-77^ Moreover, neural network biases are observed when LC activation reconfigures brain-wide functional connectivity.^45,48,78^ These findings point to a pivotal question: how do LC-NE neurons organize to modulate diverse brain functions, regardless of in a homogenous or module-like mode. It calls for multidimensional characterizations that integrate molecular, morphology, physiology, and function assays at both single-neuron and population levels.

The larval zebrafish presents an appropriate model for investigating this question due to its relatively simple and evolutionarily conserved LC system, which comprises less than 10 LC-NE neurons per hemisphere at 5 - 7 days post-fertilization (dpf), when the larva exhibits various sensorimotor functions and behaviors.^79-83^ In the present study, we integrated multiple *in vivo* methods and systematically examined the logic of the LC system in both its structural and functional organization in larval zebrafish. At a single-neuron level, employing genetically sparse labeling and *in vivo* imaging, we reconstructed the three-dimensional morphology of individual LC-NE neurons and revealed heterogeneous patterns of their axon projections. Combining *in vivo* whole-cell recording and single-cell RNA sequencing (scRNA-seq), we then found that morphologically heterogenous LC-NE neurons are comparable in electrophysiological properties, sensory-evoked responses, and expression profiles of neural function-related genes. At the population level, we further found that the collective axon projection of population LC-NE neurons within the entire LC system exhibits bilateral symmetry yet brain regional difference. By integrating *in vivo* dual whole-cell recordings, Ca^2+^ imaging, and NE release imaging, we then found that bilateral LC-NE neurons fire synchronously, causing brain-wide NE release with a regionally different pattern which mirrors that of their collective axon projections. At the function level, using *in vivo* brain-wide Ca^2+^ imaging in combination of opto- and chemo-genetic manipulations, we further revealed that the LC regulates brain-wide neural dynamics with a LC-NE neuronal activity level-dependent inverted-U characteristic. This inverted-U relationship displays regional variability across the brain, which can be attributed to the regionally distinct patterns of NE release and the regionally comparable expression profile of NE receptors (NERs). Taken together, our study delineates that individual LC-NE neurons with heterogenous morphology, comparable physiological properties, similar expression of neural function-related genes, and synchronous activities, collectively work to exert brain-wide yet regionally biased modulation, reconciling the traditional view of the LC as a homogenous unit with the emerging evidence of its modular functions.

## RESULTS

### Individual LC-NE neurons exhibit heterogenous axon projections across the brain

To label LC-NE neurons in zebrafish, we utilized the knockin line *Ki(dbh:GAL4-VP16)*, in which GAL4 is specifically expressed in cells that express the dopamine β-hydroxylase (dbh),^84^ a critical enzyme in NE biosynthesis.^85^ Immunostaining verified that this line labeled all LC-NE neurons in each larva (Figures S1A-S1C; Video S1). The average total number of LC-NE neurons was 6.8 ± 0.5 (left LC) and 6.4 ± 0.3 (right LC) at 6 dpf (Figure S1C; mean ± SEM, *p* = 0.4423). For sparse labeling of single LC-NE neurons, we used 6-dpf *Ki(dbh:GAL4-VP16);Tg(UAS:Kaede);Tg(elavl3:GCaMP5G);Et(vmat2:EGFP)* larvae with only one LC-NE neuron expressing the photo-convertible fluorescence protein Kaede (Figure S1D). In those larvae, the green signals of the calcium ion indicator GCaMP5G expressed in all neurons and the enhanced green fluorescent protein (EGFP) expressed in monoamine cells including LC-NE neurons both were used as references for aligning the reconstructed morphology of individual LC-NE neurons to the brain template of 6-dpf larval zebrafish (Figure S1D). The complex morphology of individual LC-NE neurons was imaged *in vivo* after photo-conversion of Kaede from green to red by background illumination during rearing, reconstructed using the Simple Neurite Tracer in ImageJ, and then registered into the brain template (Figure S1D; see also Methods), in which brain regions are delineated for subsequent projection analyses (Table S1).^86^

We reconstructed a total of 165 LC-NE neurons (74 in the left LC and 91 in the right LC; Figures 1A, 1B, S1E and S1F; Video S2), which are over tenfold the number of all LC-NE neurons within individual larvae (see Figure S1C).^8,9^ Their somata were located at the rhombomeres 1 and 2 (R1 and R2) in the hindbrain (Figures S1E and S1F; 118 in R1, 47 in R2). Individual LC-NE neurons extended neurite processes throughout the brain, exhibiting considerable variations in the total length (left LC: 14,625 ± 432 μm; right LC: 14,447 ± 292 μm; mean ± SEM), branch number (left LC: 301.7 ± 11.5; right LC: 303.2 ± 9.4), and process tip number (left LC: 152.4 ± 5.8; right LC: 153.1 ± 4.7), yet no significant difference was observed between the two hemispheres (Figures S1G-S1I). Moreover, these parameters followed normal distribution and showed no significant differences between left and right LC (Figures S1J-S1L), suggesting the overall similar morphological complexity between LC-NE neurons in the two hemispheres.

**Figure 1.**
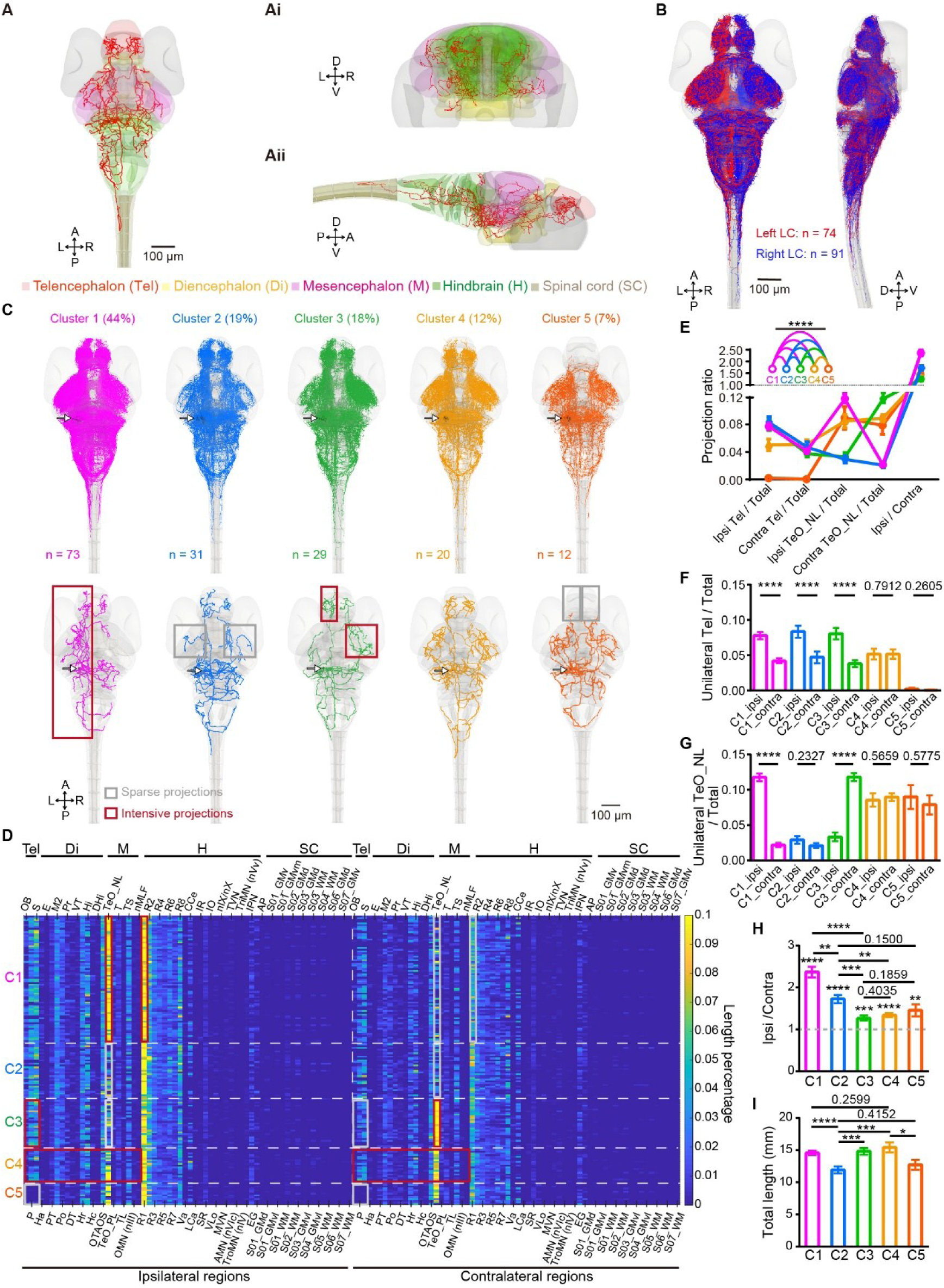
Individual LC-NE neurons exhibit brain-wide projections with heterogeneous patterns. (**A** and **B**) Reconstruction of individual LC-NE neurons’ morphology. Data were obtained from *Ki(dbh:GAL4-VP16);Tg(UAS:Kaede);Tg(elavl3:GCaMP5);Et(vmat2:EGFP)* larvae at 6 dpf, in each of which Kaede was expressed in only a single LC-NE neuron for morphology reconstruction, and the signals of GCaMP5 expressed in all neurons and EGFP expressed in monoamine neurons were used to register the reconstructed morphology of Kaede-expressing neurons to a brain template. A, anterior; P, posterior; L, left; R, right; D, dorsal; V, ventral. (**A**) Representative of brain-wide projections of one left LC-NE neuron registered in the brain template. The template encompasses five main brain areas: telencephalon (Tel), diencephalon (Di), mesencephalon (M), hindbrain (H), and spinal cord (SC), with each shown in a different color. (A) Dorsal view, (Ai) Coronal view, (Aii) Lateral view. (**B**) Collective projections of all reconstructed 74 left (red) and 91 right (blue) LC-NE neurons, shown in dorsal (left) and lateral (right) views in the brain template. The morphology of each neuron was obtained from an individual larva. (**C-I**) Characterization of the morphological features of LC-NE neurons. (**C**) Five clusters of LC-NE neurons revealed by NBLAST clustering. Top, all neurons in each cluster; Bottom, a representative neuron for each cluster. Gray rectangles, regions with sparse projections; red rectangles, regions with intensive projection; arrows, location of the somata of LC-NE neurons. (**D**) Heatmap of projection length percentages in target brain regions (length in each region/total projection length per neuron). The heatmap displays brain regions that receive projections from at least one LC-NE neuron among the all reconstructed neurons. In total, 128 brain regions, including both left and right hemispheres, spanning from the Tel to SC, were identified as targets of LC-NE neuronal projections. Tel, telencephalon; Di, diencephalon; M, mesencephalon; H, hindbrain; SC, spinal cord. All abbreviations for brain regions in Table S1. Gray rectangles, regions with sparse projections; red rectangles, regions with intensive projection. Each row represents one LC-NE neuron. C1 - C5, cluster 1 to cluster 5. (**E**) Projection ratio for C1 - C5, calculated as the projection length within target regions divided by the total projection length across all regions, or calculated as ipsilateral projection length divided by contralateral projection length (only for the rightmost data points). Ipsi, ipsilateral; contra, contralateral. Tel, telencephalon; TeO_NL, optic tectum neuropil layer. One-way multivariate analysis of variance (MANOVA) was performed for pairwise comparisons between clusters. (**F** and **G**) Projection ratio of unilateral Tel (**F**) or unilateral TeO_NL (**G**) for C1 - C5, calculated as the projection length within unilateral target region divided by the total projection length across all brain regions. Paired t-test was used for intra-cluster comparisons. (**H**) Ratio of ipsilateral to contralateral projection lengths for C1 - C5. For comparison with 1 for each cluster, C1 - C4 conform to a normal distribution (Kolmogorov-Smirnov test) and were analyzed using a one-sample t test. C5 does not conform to a normal distribution and was analyzed using a one-sample Wilcoxon signed-rank test. Unpaired t-test was applied for inter-cluster comparisons. (**I**) Averaged total projection length per neuron in each cluster. Unpaired t-test was performed. **\****p* < 0.05, **\*\****p* < 0.01, **\*\*\****p* < 0.001, **\*\*\*\****p* < 0.0001. Data are represented as mean ± SEM. See also Figure S1, Videos S1 and S2, and Table S1.

To discriminate the axon and dendrite within the neurite processes of LC-NE neurons, we used *Ki(dbh:GAL4-VP16);Tg(UAS:mCherry);Tg(UAS:sypb-EGFP)* larvae, in which axonal varicosities of LC-NE neurons were labeled by the synaptic vesicle protein synaptophysin b (sypb) fused with EGFP.^87^ We found that LC-NE neuronal dendrites, which lacked sypb-EGFP expression, were mainly localized in ipsilateral R1 and R2, and exhibited relatively short processes (Figures S1M and S1N; see Methods). Based on the dendritic properties, we manually classified dendrites and axons of all the 165 reconstructed LC-NE neurons. The dendrites, of which 58% distributed in R1 and 31% in R2, accounted for less than 7% of the total neurite length (Figures S1O-S1Q; left LC: 972.1 ± 42.7 μm for dendrites, 13,653 ± 410.0 μm for axons, 0.067 ± 0.0024 for dendrite/axon length ratio; right LC: 932.8 ± 31.8 μm for dendrites, 13,514 ± 276.7 μm for axons, 0.065 ± 0.0018 for dendrite/axon length ratio).

Then we mirrored all right LC-NE neurons to the left hemisphere along the midline of the brain template and performed NBLAST clustering of the morphology of 165 reconstructed LC-NE neurons.^88^ Five clusters were identified (named as C1, C2, C3, C4 and C5), and each of them exhibited distinct projection patterns across the brain with brain region bias and lateral preference (Figures 1C-1I). Notably, the ipsilateral projections in R1 and R2 (Figure 1D) are mainly due to the dendrites of LC-NE neurons. C1 neurons (44%, n = 73) displayed strong ipsilateral projection preference. C2 neurons (19%, n = 31) had fewer arborizations in bilateral optic tectum neuropil layers (TeO_NL). C3 neurons (18%, n = 29) showed ipsilateral projection preference in the telencephalon (Tel) and contralateral preference in TeO_NL. Both C4 (12%, n = 20) and C5 neurons (7%, n = 12) showed relatively balanced bilateral projections, but C5 neurons had rare arborizations in Tel (Figures 1E-1H). In addition, the somata of the five clusters’ neurons were intermingled within the regions R1 and R2 (Figure S1R). Taken together, these results reveal the morphological heterogeneity of LC-NE neurons.

### Individual LC-NE neurons display comparable physiological properties

To investigate whether physiological properties differ among different morphology clusters, we performed *in vivo* whole-cell recording on morphologically identified LC-NE neurons, and then sequentially characterized their intrinsic electrophysiological properties and sensory-evoked responses (n = 35; Figures S2A and 2A). Among 35 LC-NE neurons examined, 9 cells were intentionally collected outside the 165 LC-NE neurons analyzed above, because during random electrophysiological recording, neurons belonged to some clusters (e.g., C4 and C5) were rarely encountered due to their low percentages (Figure 2B). To preserve the randomness of the sampling during morphology reconstruction, these 9 neurons were excluded from the morphological clustering analysis.

**Figure 2.**
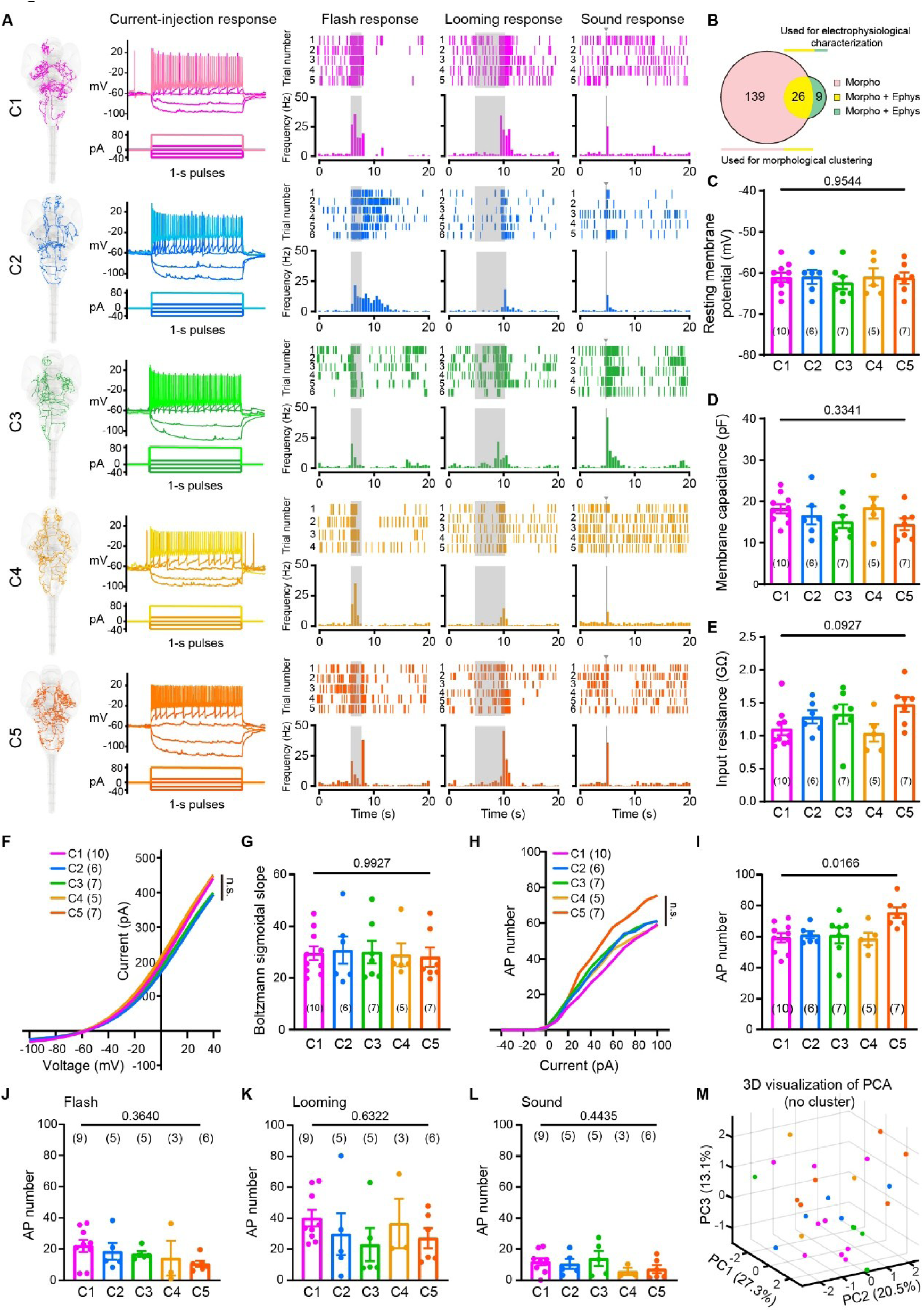
Individual LC-NE neurons display comparable physiological properties. (**A**) Representative cases of whole-cell recordings from C1 - C5. For each case, morphology (left), current injection-evoked responses (middle), and raster plots and the frequency histograms of sensory responses (right) were shown. Grey shadows indicate the time window of sensory stimuli (2-s flash, 5-s looming, and 10-ms pure tone). (**B-I**) Summary of intrinsic electrophysiological properties of LC-NE neurons in C1 - C5. The number in the brackets represents the number of recorded LC-NE neurons in each cluster in (**C-L**). (**B**) Composition of datasets used for electrophysiological characterization as well as morphological clustering. Of the 35 LC-NE neurons characterized electrophysiologically, 26 were also part of the 165 neurons used for morphological clustering, while the remaining 9 were intentionally collected and excluded from morphological clustering. Ephys, electrophysiological recording; Morpho, morphology reconstruction. (**C-E**) Summary of the resting membrane potential (**C**), membrane capacitance (**D**), and input resistance (**E**) of LC-NE neurons in C1 - C5. (**F**) I-V curves of LC-NE neurons in C1 - C5. Colored lines represent the average curves for all LC-NE neurons within each morphological cluster, fitted using Boltzmann’s equation. (**G**) Boltzmann sigmoidal slope of LC-NE neurons in C1-C5, referring to the slope of the curve fitted with Boltzmann’s equation. (**H**) Number of APs evoked by current injections of LC-NE neurons in C1 - C5. Current injections are from -40 pA to 100 pA with a 10-pA step lasting 1 s. Colored lines represent the average curves for all LC-NE neurons within each cluster. (**I**) Number of APs evoked by 100-pA current injections of LC-NE neurons in C1 - C5. One-way ANOVA test shows significant difference among clusters, and further post-hoc analysis using Tukey’s multiple comparisons test reveals significant differences only between C1 and C5 (*p* = 0.0171) and between C4 and C5 (*p* = 0.0452), with no significant difference between other group pairs. (**J-L**) Number of APs evoked by flash (**J**), looming (**K**), and pure tone (**L**) in LC-NE neurons of C1 - C5. Data were obtained from 28 LC-NE neurons, out of the total 35 used for analyzing intrinsic electrophysiological properties. Each data point represents the average value of 4 - 10 trials for each neuron. (**M**) 3D visualization of principal component analysis (PCA) of the collective properties of 28 LC-NE neurons based on 11 parameters, including the resting membrane potential, membrane capacitance, input resistance, Boltzmann sigmoidal slope of I-V curve, number of APs evoked by a 100-pA current injection, number of APs evoked by flash, looming, and pure tone stimuli (see **C-E**, **G**, **I-L**), and the corresponding firing frequencies for each sensory modality (see Figures S2C-S2E). The optimal number of clusters was determined to be 1 based on the gap statistic analysis (see Figure S2F and Methods). Each point in (**C-E**, **G**, **I-M**) represents the data obtained from an individual neuron. One-way ANOVA test was performed (**C-E**, **G**, **I-L**). Two-sample Kolmogorov-Smirnov test was performed between different morphological clusters (**F**, **H**). Data are represented as mean ± SEM. See also Figure S2.

Whole-cell recording revealed that the physiological properties of LC-NE neurons were comparable within and across C1 - C5 neurons (Figures 2 and S2). Statistical analyses showed no significant overall difference in intrinsic electrophysiological properties across C1 - C5 (Figures 2C-2I), including the resting membrane potential (- 61.3 ± 0.6 mV), membrane capacitance (16.1 ± 0.7 pF), input resistance (1.2 ± 0.1 GΩ), current-voltage (I-V) curve, and current injection-evoked firing pattern. Post-hoc analysis showed that the only significant difference was in the number of action potentials (APs) evoked by current injections, observed between C1 and C5 (*p* = 0.0171) and between C4 and C5 (*p* = 0.0452) (Figure 2I). Subsequent characterizations of the sensory responses of 28 recorded neurons evoked by visual (flash and looming) and auditory (pure tone) stimuli showed that the same stimuli evoked comparable firing patterns, including the number and frequency of APs, in neurons across different clusters (Figures 2A, 2J-2L, and S2B-S2E). Furthermore, based on these five intrinsic electrophysiological properties and six properties of responses evoked by three types of sensory stimuli, i.e. 11 parameters in total, we carried out the gap statistic analysis of all the 28 neurons examined and found no obvious cluster (Figures 2M, S2F and S2G).^89^ Taken together, these results suggest that individual LC-NE neurons, while exhibiting heterogeneous axon projections, possess largely comparable physiological properties.

### Individual LC-NE neurons show comparable expression patterns of neural function-related genes

To investigate the diversity of LC-NE neurons at the molecular level, we performed scRNA-seq to analyze the transcriptomic profiles of LC-NE neurons. Using 6-dpf *Ki(dbh:GAL4-VP16);Tg(4×nrUAS:GFP)* larvae, we isolated GFP-expressing LC-NE neurons for single-cell SMART sequencing (Figure 3A; see Methods).^90^ In total, we successfully sequenced 209 eligible LC-NE neurons, achieving an average library mapping rate of 87.1%, a median of 7,200 genes detected per cell, and a mitochondrial gene content median of 0.9% (Figures S3A-S3D). The marker genes of LC-NE neurons, including *dbh, th,* and *slc6a2*, confirmed the cell specificity (Figure S3E). Based on the top 2,000 variable genes identified through principal component analysis (PCA), LC-NE neurons were classified into three clusters (Figure 3B). Gene Ontology (GO) analysis of the top 20 marker genes for each cluster (Figure S3F) revealed that these markers were primarily associated with fundamental biological functions (Figure 3C), including ATP-dependent protein folding chaperones, heat shock protein binding, and transcription factor binding, etc.

**Figure 3.**
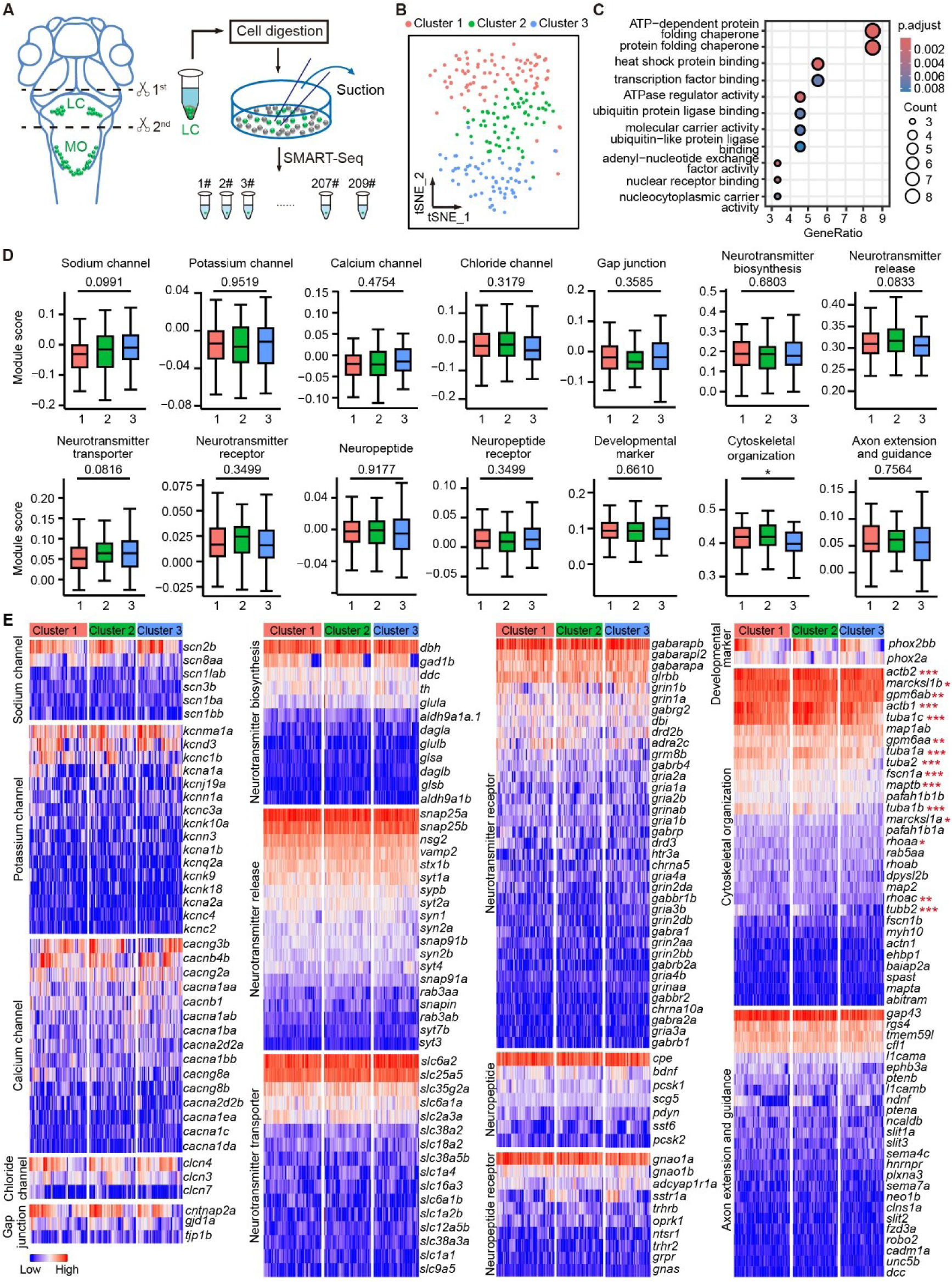
Individual LC-NE neurons show comparable expression profiles of neural function-related genes. (**A**) Schematic of anatomical sampling of brain tissue containing LC-NE neurons, followed by the collection of GFP-expressing LC-NE neurons and subsequent SMART-Seq. The sampling was performed in *Ki(dbh:GAL4-VP16);Tg(4×nrUAS:GFP)* larvae at 6 dpf. A total number of 209 LC-NE neurons were obtained in four times experiments for single-cell sequencing. MO, medulla oblongata. (**B**) t-distributed stochastic neighbor embedding (tSNE) plot, showing 3 clusters of 209 LC-NE neurons using Seurat analysis. In total, cluster1 contains 81 cells, cluster2 contains 65 cells, and cluster3 contains 63 cells. Each cluster is color-coded. (**C**) Gene ontology (GO) analysis of top 20 marker genes (listed in Figure S3F) of 3 clusters in (**B**). (**D**) Boxplots showing the calculation of each gene category activity among 3 clusters using the area under curve (AUC) score. Genes of total 14 categories are listed in Table S2. One-way ANOVA test was performed. (**E**) Heatmap showing the expression level of the total of 202 genes among 14 gene categories in 3 clusters. All these genes were expressed in above 30% LC-NE neurons. The red asterisks indicate significance among 3 clusters. **\****p* < 0.05, **\*\****p* < 0.01, **\*\*\****p* < 0.001. Data are represented as mean ± SEM. See also Figure S3 and Table S2.

To further explore the expression of morphological development- and physiology-related genes across the three clusters, we analyzed 14 key categories of genes, encompassing 721 genes, among which 654 were expressed in zebrafish LC-NE neurons (Table S2). These categories included developmental markers and genes associated with morphological growth (including cytoskeletal organization, and axon extension and guidance), neuronal excitability (including membrane ion channels for transporting sodium, potassium, calcium, and chloride ions, and gap junctions), neurotransmitter processing (including biosynthesis, release, and transportation), and neurotransmitter receptors (including excitatory, inhibitory, and neuropeptides). Genes associated with neural functions were expressed with no significant difference across all three clusters (Figure 3D), providing a potential basis for the comparable physiological properties observed among individual LC-NE neurons above. Notably, genes associated with morphological development exhibited differential expression, particularly those related to cytoskeletal organization, such as *tuba1a*, *tuba1b*, *tuba1c*, *tuba2*, *maptb*, *actb1*, and *actb2* (Figures 3D and S3G), aligning with the morphological heterogeneity observed within LC-NE neurons. Similar results were observed by heatmap analysis of these category genes expressed in over 30% of LC-NE neurons (Figure 3E). These results indicate the comparable expression of neural function-related genes across LC-NE neurons, along with differential expression of development-related genes.

### Population LC-NE neurons display bilaterally symmetrical yet regionally distinct axon projections

To investigate the anatomical organization of the entire LC system, we then examined the projection pattern of population LC-NE neurons across the brain. As the number of neurons in the bilateral LCs of individual larvae is relatively equal (see Figure S1C), we analyzed the collective axon projections of all 74 reconstructed left LC-NE neurons and a randomly sampled subset of 74 cells from the 91 reconstructed right LC-NE neurons. Both left and right LC-NE neurons exhibited a similar preference for ipsilateral projections (Figures 4A and 4B; see also Figure 1H). Then we quantified the average axon projections in each brain region, and found that both the axon length and density were comparable in bilateral regions but showed significant differences along the anterior-posterior axis (Figures 4C, 4D, S4A and S4B), with the telencephalon (Tel) and hindbrain (H) regions exhibiting a higher axon density compared to the diencephalon (Di) and midbrain (M) (Figure 4D). Similar results were achieved when the entire LC-NE system in an individual larva was simulated by randomly sampling 7 left and 7 right LC-NE neurons per group from the 165 reconstructed neurons, in accordance with the proportions of the cluster composition (Figures 4E-4G, S4C and S4D; Video S2). It should be mentioned that LC-NE neuronal dendrites contribute to the distribution within R1 and R2 (see Figure 1D).

**Figure 4.**
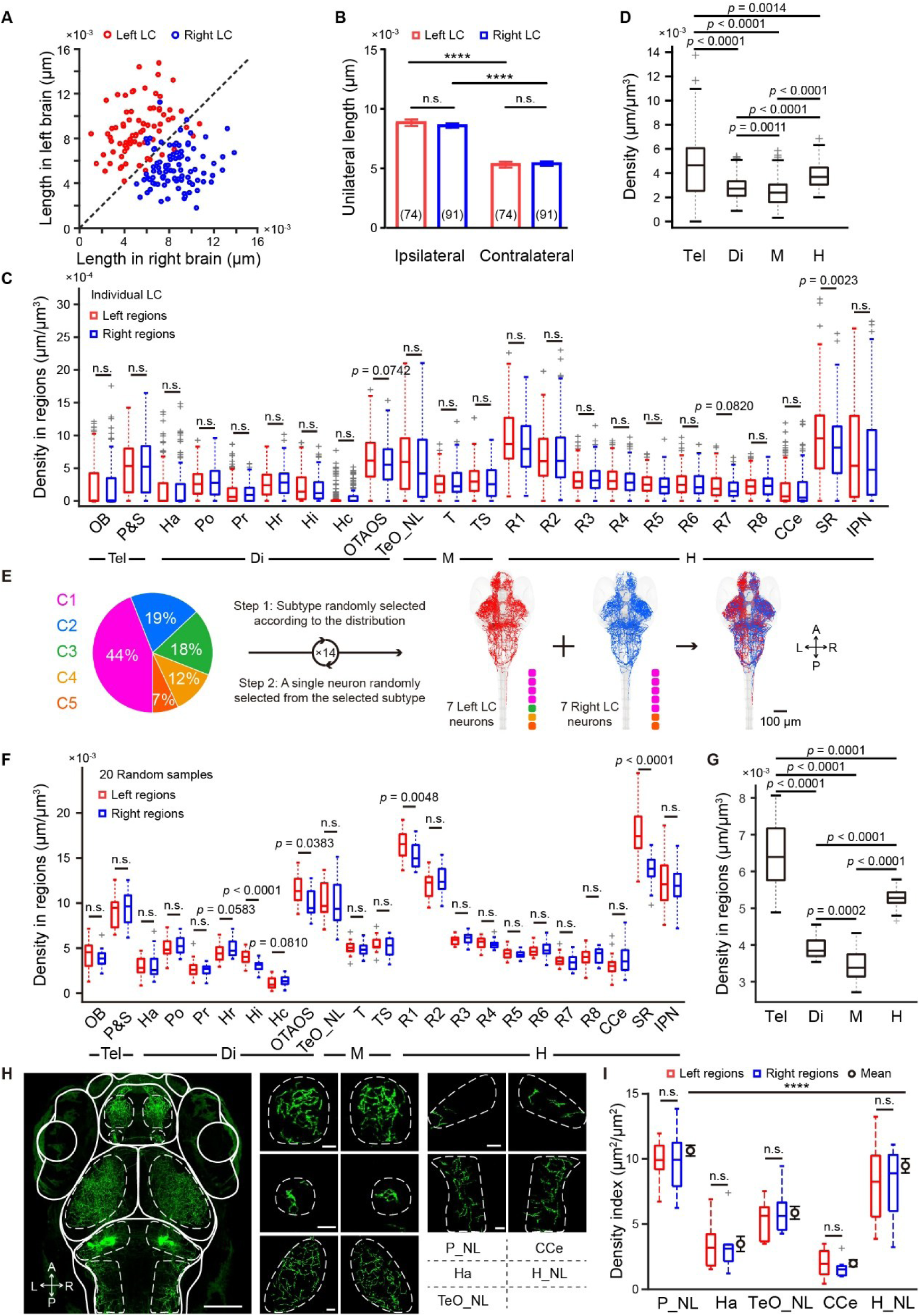
Population LC-NE neurons display bilaterally symmetrical yet regionally distinct axon projections. (**A-D**) Projection distribution of collective LC-NE neurons reconstructed. The dataset is the same as that in Figure 1. (**A**) Projection lengths of individual neurons in both left and right hemispheres. Red dots, left LC-NE neurons; blue dots, right LC-NE neurons. The dashed line is diagonal. (**B**) Average lengths of ipsilateral and contralateral projections of left (red) and right (blue) LC-NE neurons. Paired t-test was performed for comparisons between ipsilateral and contralateral projections of unilateral LC-NE neurons, while unpaired t-test was performed for comparisons between bilateral LC-NE neurons. (**C**) Average density of LC-NE neuronal projections in each brain region of the left (red) and right (blue) hemispheres. This was calculated as the average density of all 74 left LC-NE neurons and 74 randomly selected neurons from 91 right LC-NE neurons. The median was represented by a central line, while the interquartile range (IQR) was defined by the first and third quartiles, displayed as box edges. The whiskers were extended to the most extreme non-outlier data points, and values exceeding this range were marked as outliers (grey markers). Tel, telencephalon; Di, diencephalon; M, mesencephalon; H, hindbrain. (**D**) Average projection density of all 165 LC-NE neurons in brain areas along the anterior-posterior axis. (**E-G**) Simulation of the entire LC system in an individual larva by cluster-based proportional random sampling from 165 individual LC-NE neurons reconstructed. (**E**) Schematic of cluster-based proportional random sampling. According to percentages of 5 morphological clusters, 7 left (red) and 7 right (blue) LC-NE neurons were randomly selected to simulate the entire LC system in an individual larva. Random sampling was performed 20 times for statistical analysis. (**F**) Projection densities of the simulated entire LC system in each brain region of the left (red) and right (blue) hemispheres. (**G**) Projection densities of the simulated entire LC system in brain areas along the anterior-posterior axis. (**H** and **I**) Projections of the real LC system. Data were obtained from six *Ki(dbh:GAL4-VP16);Tg(4×nrUAS:GFP)* larvae at 6 dpf. P_NL, pallium neuropil layer; Ha, habenula; TeO_NL, optic tectum neuropil layer; CCe, cerebellum corpus cerebelli; H_NL, hindbrain neuropil layer. (**H**) Representative of projected confocal images showing the LC system in a 6-dpf larva, in which 7 left and 7 right LC-NE neurons are labeled. Brain regions indicated by dashed lines are enlarged (right). (**I**) Projection density in brain regions along the anterior-posterior axis. The middle section of each brain region was selected for analysis. Fluorescent images were auto-binarized using Fiji software, and region of interest (ROIs) were delineated. The ratio of fluorescent-positive pixels to the total area was calculated as the density index for the corresponding ROI using the Fiji plugin ROI manager. Wilcoxon matched-pairs signed-rank test was performed for comparison between counterpart bilateral regions, while one-way ANOVA test was performed for comparison among different regions. For two-group comparisons, a paired t-test was used for normally distributed data, while the Wilcoxon matched-pairs signed-rank test was applied for non-normally distributed data (**C**, **F**). Unpaired t-test was applied for inter-regional comparisons (**D, G**). n.s., not significant; **\*\*\*\****p* < 0.0001. See also Figure S4 and Video S2-S4.

Furthermore, we mapped the collective processes of all LC-NE neurons in individual *Ki(dbh:GAL4-VP16);Tg(4×nrUAS:GFP)* larvae and analyzed their density in the pallium (P) of the telencephalon, habenula (Ha) in the diencephalon, optic tectum (TeO) in the midbrain, cerebellum corpus cerebelli (CCe), and the hindbrain (Figure 4H; Video S3), where LC-NE neuron axons were densely distributed. As the axons in the pallium, optic tectum, and hindbrain predominantly distributed in neuropil layers (NLs), the analysis in these regions was focused on their NLs (i.e., P_NL, TeO_NL, and H_NL). As the neurite processes of all LC-NE neurons in individual larvae were too dense to reconstruct, the fluorescence pixel density was used as an index for quantification. Consistently, the axon projections showed bilateral symmetry but significant diversity among different regions (Figure 4I). To exclude the projections of medulla oblongata noradrenergic (MO-NE) neurons, we conducted two-photon laser-based ablation of bilateral LCs in *Ki(dbh:GAL4-VP16);(Tg(4×nrUAS:GFP)* larvae and examined exclusive axon projections of MO-NE neurons. The result showed that MO-NE projections were confined primarily to the ventral diencephalon, ventral hindbrain, medulla, and spinal cord, and were spatially segregated from LC-NE projections (Video S4).

These results suggest that the axon projections of the LC-NE system are bilaterally symmetrical but regionally distinct, possibly causing differential NE release and functional modulation across brain regions.

### Population LC-NE neurons exhibit synchronous spontaneous activities and uniform sensory responses

To link the LC axon projection patterns with potential downstream effects, we investigated the activity of population LC-NE neurons within intact zebrafish. Using *in vivo* dual whole-cell recordings of paired LC-NE neurons in either ipsilateral or contralateral hemispheres, we observed that they exhibited both spontaneous tonic and phasic firing patterns (Figures 5A and 5B; Video S5), aligning with previous findings in rodents and non-human primates.^23,43,46,47^ To quantify the firing synchrony, we assessed the time delay between APs of the paired LC-NE neurons (Figures 5B and 5C). LC-NE neurons displayed highly synchronized phasic firing, characterized by a minimal delay (unilateral pairs: 7.2 ± 0.7 ms; bilateral pairs: 6.9 ± 0.5 ms; *p* = 0.4143). In contrast, tonic firing was associated with significantly longer delays (*p* < 0.0001) (unilateral pairs: 130.2 ± 16.2 ms; bilateral pairs: 110.4 ± 12.9 ms; *p* = 0.9461). Furthermore, we conducted *in vivo* Ca^2+^ imaging in *Ki(dbh:GAL4-VP16);Tg(UAS:GCaMP6s)* larvae, in which the calcium indicator GCaMP6s was selectively expressed in LC-NE neurons, and observed that bilateral populations of LC-NE neurons showed synchronous spontaneous Ca^2+^ activities (Figures 5D-5G).

**Figure 5.**
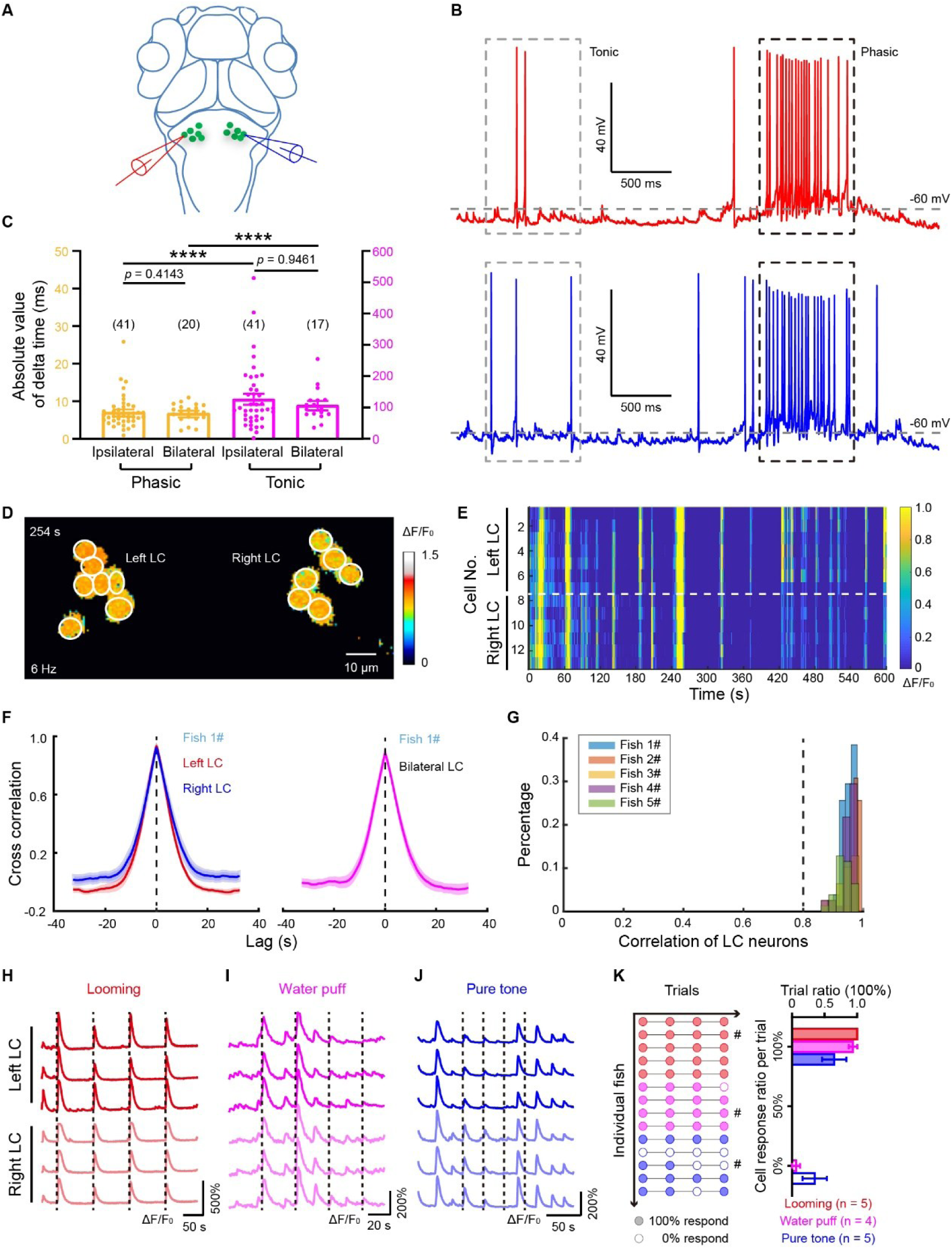
Population LC-NE neurons exhibit synchronous spontaneous activities and uniform sensory responses. (**A-C**) Synchronous spontaneous activities of LC-NE neurons examined by *in vivo* dual whole-cell recordings. (**A**) Schematic of *in vivo* dual whole-cell recordings of a pair of bilateral LC-NE neurons in *Ki(dbh:GAL4-VP16);Tg(4×nrUAS:GFP)* larvae at 6 dpf. (**B**) Representative recording traces of a pair of bilateral LC-NE neurons recorded, showing synchronous spontaneous activities between the two neurons. Activities were manually segmented into tonic (gray dashed box) and phasic (black dashed box) events for calculation of time delays between APs of the paired LC-NE neurons in (**C**). (**C**) Averaged time delay between two temporally closest APs (i.e., paired APs) of simultaneously recorded LC-NE neurons. The time delay of paired APs is calculated as the time difference between the two APs. In total, 41 phasic and 41 tonic events from 7 unilateral pairs, and 20 phasic and 17 tonic events from 3 bilateral pairs were analyzed. (**D-G**) Synchronous spontaneous activities of population LC-NE neurons examined by *in vivo* confocal calcium imaging. (**D-F**) Synchronous spontaneous calcium activities of the somata of 7 left and 6 right LC-NE neurons in a *Ki(dbh:GAL4-VP16);Tg(UAS:GCaMP6s)* larva at 6 dpf. Imaging rate, 6 Hz. (**D**) Color-coded ΔF/F_0_ of calcium activities at the time point of 254 second in (**E**). (**E**) Heatmap of the spontaneous calcium activities of all the 13 LC-NE neurons. Left (up) and right (down) LC-NE neurons are separated by a dashed line. (**F**) Averaged pairwise cross correlation of the spontaneous activities of population LC-NE neurons shown in (**E**). Left: unilateral pairs of LC-NE neurons. Right: bilateral pairs of LC-NE neurons. Shaded error bars represent SEM. (**G**) Distribution of the cross correlation among all pairs of LC-NE neurons within five individual larvae. Each color represents data obtained from one larva at 6 dpf. (**H-K**) Uniform sensory responses of LC-NE neurons examined by *in vivo* confocal calcium imaging. (**H-J**) Representative calcium activities evoked by visual looming (**H**), water puff at the tail (**I**), and pure tone (J), with each trace representing a single LC-NE neuron imaged from 6-dpf *Ki(dbh:GAL4-VP16);Tg(UAS:GCaMP6s)* larvae at a single optic plane. The data in (**I** and **J**) were obtained from the same larva. Dashed lines indicate the onset of the sensory stimuli. (**K**) All-or-none responsiveness of LC-NE neurons. Left: responsiveness of LC-NE neurons for each trial in individual larvae. Filled circles, trials with all LC-NE neurons responsive; open circles, trials with all LC-NE neurons unresponsive. The number indicates individual larvae examined. Note: some larvae were tested using multiple types of sensory stimuli. Right: trial ratio of fully responsive or fully unresponsive. #, representative responses shown in (**H-J**). Two-tailed Mann-Whitney test was performed (**C**). **\*\*\*\****p* < 0.0001. Data are represented as mean ± SEM. See also Figure S5 and Video S5.

We further examined their sensory responses evoked by visual (looming), mechanical (water puff at the tail), and auditory (pure tone) stimuli. In the majority of trials, all LC-NE neurons displayed robust responses to these stimuli with phasic activities (Figures 5H-5K; see also Figures 2 and S2). Notably, the entire population of bilateral LC-NE neurons exhibited uniform sensory responsiveness in an “all-or-none” mode, i.e. all responsive or all non-responsive (Figures 5H-5K).

The activity synchrony of bilateral LC-NE neurons suggests that these neurons may receive common synaptic inputs and/or electrically couple together. Prior studies reported that gap junctions involve in synchronous activities of LC-NE neurons.^23,35,91^ The single-cell SMART sequencing data showed expression of gap junction-associated genes, including *cntnap2a* and *gjd1a*, in LC-NE neurons (see Figure 3E). We then loaded neurobiotin into recorded LC-NE neurons via whole-cell micropipettes (see Methods)^92^ and observed that neurobiotin immunostaining was prominent in most ipsilateral LC-NE neurons but rarely observed in contralateral LC-NE neurons. (Figures S5A-S5C; ipsilateral *vs* contralateral: 91.2 ± 3.1% *vs* 10.0 ± 5.1%, *p* < 0.001). Furthermore, we performed dual whole-cell recordings of LC-NE neurons and found that ipsilateral LC-NE neurons showed electrical coupling, whereas contralateral neurons did not (Figures S5D-S5H; 5/5 pairs for ipsilateral, 0/7 pairs for contralateral).

Collectively, these results indicate that the activities of population LC-NE neurons are highly synchronous and uniform, a characteristic likely attributable to gap junctions and/or common synaptic inputs for ipsilateral neurons, and common inputs for bilateral neurons.

### Brain-wide NE release displays globally synchronized and regionally distinct patterns

The bilaterally symmetrical yet regionally distinct axon projections of population LC-NE neurons, along with their synchronized activities, implies regionally patterned NE release across the brain. To investigate this point, we performed *in vivo* time-lapse imaging in 6-dpf *Tg(elavl3:GRAB_NE1h_)* larvae, in which the G-protein-coupled receptor (GPCR) activation-based NE sensor GRAB_NE1h_ is driven by the pan-neuronal *elavl3* promoter to express in neurons for detecting NE release.^40^ Spontaneous NE release was detected across the brain, exhibiting synchrony in various regions including P_NL, Ha, TeO_NL, CCe and H_NL (Figures 6A and 6B), where LC-NE neuron axons were densely distributed (see Figure 4H). The peak amplitudes of NE release in bilateral regions were comparable (Figure 6C, scattered groups), reflecting the bilateral symmetry of population LC-NE neurons’ projections (see Figures 4 and S4). Notably, significant differences in the amplitude were observed along the anterior-posterior axis (Figure 6C, average groups, *p* < 0.0001), consistent with the distribution of population LC-NE neurons’ projection densities (see Figure 4I). In line with this, the peak amplitude of NE release was found to positively correlate with the projection density of LC-NE neurons across the brain (Figure 6D, *p* = 0.0002), emphasizing the significance of the structure in determining the strength of NE release.

**Figure 6.**
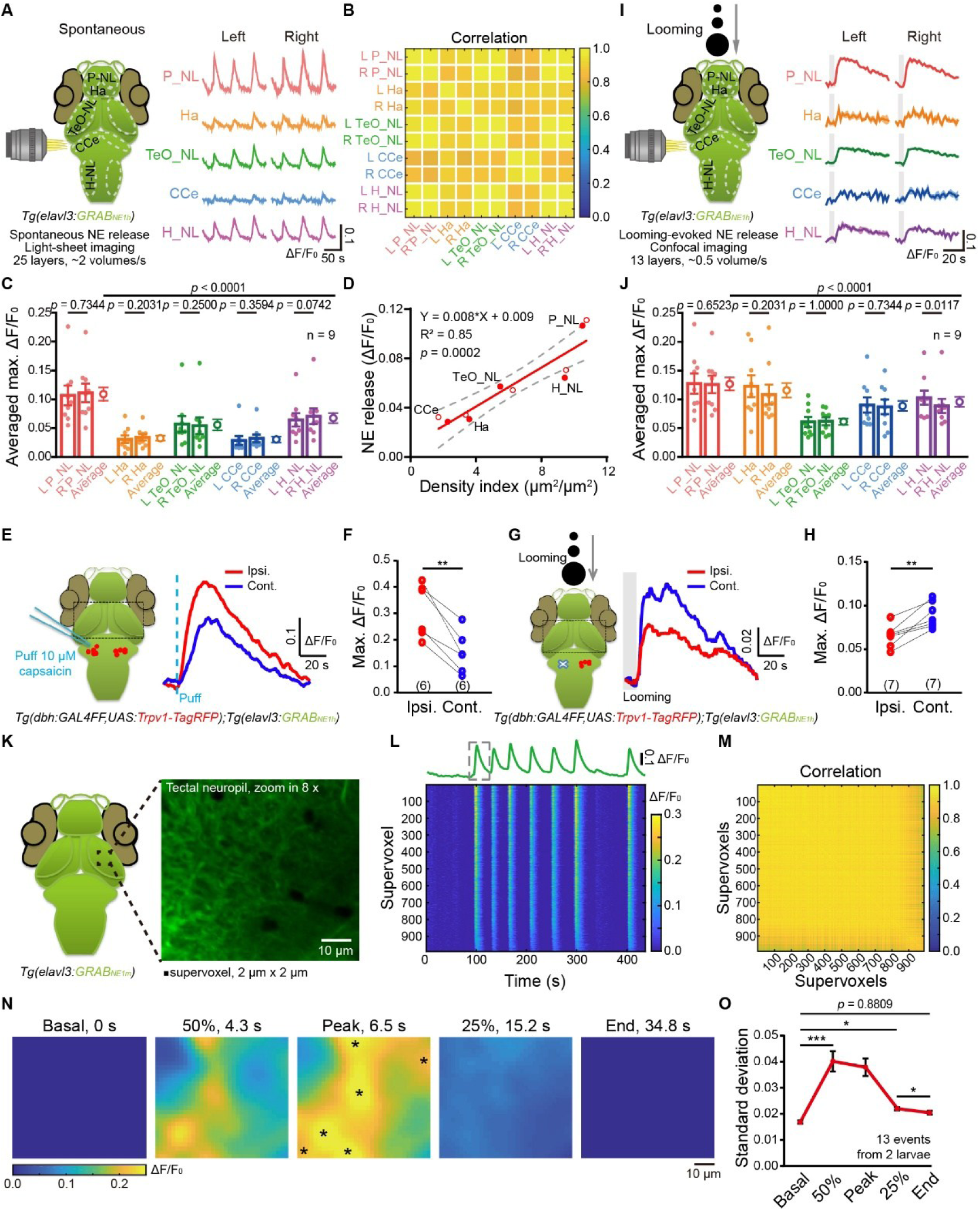
Brain-wide NE release displays globally synchronized and regionally distinct patterns. (**A-D**) Brain-wide synchronous spontaneous NE release revealed by *in vivo* light-sheet imaging of *Tg(elavl3:GRAB_NE1h_)* larvae at 6 dpf. P_NL, pallium neuropil layer; Ha, habenula; TeO_NL, optic tectum neuropil layer; CCe, corpus cerebelli; H_NL, hindbrain neuropil layer. (**A**) Left: schematic of experiments. Background green signals indicate brain-wide expression of GRAB_NE1h_ sensors in neurons. The white dashed lines indicate the analyzed brain regions. Right, typical traces of spontaneous NE release in bilateral regions. (**B**) Correlation of spontaneous NE release among brain regions. L-P_NL, right P_NL; R-P_NL, right P_NL; L-Ha, left Ha; R-Ha, right Ha; L-TeO_NL, left TeO_NL; R-TeO_NL, right TeO_NL; L-CCe, left CCe; R-CCe, right CCe; L-H_NL, left H_NL; R-H_NL, right H_NL. (**C**) Peak amplitude of NE release in different brain regions. Data were obtained from 9 events from 5 larvae. Each dot represents one event. Mean peak amplitude of counterpart regions was listed at right. (**D**) Positive correlation between the peak of NE release and the axon projection density of LC-NE neurons. The data were fitted with least-squares linear regression (red line). Dotted lines show the 95% confidence bands of the best-fit line. (**E-H**) Effects of unilateral manipulations of LC-NE neurons on evoked NE release, revealed by *in vivo* confocal imaging of *Tg(dbh:GAL4FF,UAS:TRPV1-TagRFP);Tg(elavl3:GRAB_NE1h_)* larvae at 6 dpf. (**E**) NE release evoked by the chemogenetic activation of unilateral LC-NE neurons via local puffing of 10 μM capsaicin. The black dashed box indicates the TeO_NL region of imaging. Right, NE release traces of bilateral TeO_NL. The dash cyan line indicates the onset of the local puffing. Ipsi., ipsilateral TeO_NL, in magenta; Cont., contralateral TeO_NL, in blue. (**F**) Peak amplitude of chemogenetic activation-evoked NE release in ipsilateral and contralateral TeO_NL. Data were obtained from 6 larvae. (**G**) NE release evoked by looming after two-photon laser-based unilateral LC ablation in *Tg(dbh:GAL4FF,UAS:Trpv1-TagRFP)* larvae. The black dashed box indicates the TeO_NL region of imaging. Right, NE release traces of bilateral TeO_NL. The grey shadow indicates the time window of looming. Ipsi., ipsilateral TeO_NL, in magenta; Cont., contralateral TeO_NL, in blue. (**H**) Peak amplitude of looming-evoked NE release in ipsilateral and contralateral TeO_NL. Data were obtained from 7 larvae. (**I** and **J**) Looming-evoked brain-wide NE release revealed by *in vivo* confocal imaging of *Tg(elavl3:GRAB_NE1h_)* larva at 6 dpf. (**I**) Left: schematic of experiments. Right, trial-averaged traces of looming-evoked NE release in bilateral regions. Looming stimuli lasting for 5 s were presented 5 times with a 90-s interval. (**J**) Peak amplitude of looming-evoked NE release in different brain regions. Data were obtained from 9 larvae. Each dot represents one event. Mean peak amplitude of counterpart regions was listed at right. (**K-O**) Fine-scale characterization of local spontaneous NE release revealed by *in vivo* confocal imaging of *Tg(elavl3:GRAB_NE1m_)* larvae at 6 dpf. (**K**) Schematic of experiments. The black box indicating the imaging region (0.1554 μm/pixel) of TeO_NL was shown in right. A single plane image was segmented into 2 *×* 2 μm^2^ supervoxels for analysis. (**L**) Pixel-wise averaged trace (top) and heatmap (bottom) of spontaneous NE release within the imaging region in (**K**). The gray dashed box represents one NE release event for spatiotemporal pattern analysis in (**N**). (**M**) Correlation of spontaneous NE release among supervoxels within the imaging region in (**K**). (**N**) Spatiotemporal pattern of the NE release event shown in the dashed box in (**L**). Heatmap with color-coded amplitudes of five frames are shown. They represent different stages of NE release signals: basal level (0 s), up to 50% of the peak (4.3 s), peak level (6.5 s), down to 25% of the peak (15.2 s), down to basal level (end, 34.8 s). Asterisks, hotspots of NE release. (**O**) Comparison of the standard deviations of the NE release event at different stages. 13 events from 2 larvae were analyzed. Wilcoxon signed rank test was performed for comparison between counterpart regions, and one-way ANOVA Friedman test was performed for comparison along the anterior-posterior axis (**C**, **J**). The t-test was used to test whether the slope is significantly different from 0 (**D**). Paired t-test was performed (**F**, **H**). Wilcoxon matched-pairs signed-rank test was performed (**O**). **\****p* < 0.05, **\*\****p* < 0.01, **\*\*\****p* < 0.001. Data are represented as mean ± SEM. See also Video S6.

To validate the role of LC-NE neuronal projections in NE release level, we monitored NE release evoked by chemogenetic activation of LC-NE neurons in 6-dpf *Tg(dbh:GAL4-VP16,UAS:TRPV1-TagRFP);Tg(elavl3:GRAB_NE1h_)* larvae, in which LC-NE neurons expressed the mammalian transient receptor potential vanilloid 1 (TRPV1), a ligand-gated cation channel that can be activated by capsaicin, fused with the red fluorescent protein TagRFP,^93^ and the NE sensor GRAB_NE1h_ were expressed in neurons.^40^ Chemogenetic activation of unilateral LC-NE neurons via locally puffing capsaicin to the LC soma region induced a larger NE release signal in the ipsilateral TeO_NL (Figures 6E and 6F), whereas two-photon laser-based ablation of unilateral LC significantly reduced looming-evoked NE release in the ipsilateral TeO_NL in comparison with the contralateral region (Figures 6G and 6H). These changes in NE release are consistent with ipsilateral preference of axon projections of LC-NE neurons (see Figures 4A and 4B).

We then examined the pattern of NE release in response to looming stimuli. As observed under spontaneous conditions (see Figures 6A-6C), NE release induced by looming was comparable between bilateral hemispheres (Figures 6I and 6J, scattered groups), and significant variations along the longitudinal axis were also observed (Figure 6J, average groups, *p* < 0.0001). Compared to spontaneous NE release, the overall level of looming-evoked NE release was higher, with particular increases in the Ha and CCe (compare Figures 6C and 6J). These results imply that visual inputs may locally regulate NE release within some regions, besides their direct effects on NE release via activating LC-NE neurons.

Beyond the global pattern of NE release, we further delved into the spatiotemporal dynamics of NE release at a fine scale, utilizing high-resolution imaging on *Tg(elavl3:GRAB_NE1m_)* larvae,^40^ as exemplified in the TeO_NL (Figure 6K). Spontaneous NE release exhibited marked synchronization even at the level of supervoxels (2 μm × 2 μm) (Figures 6L and 6M). Moreover, we observed discrete hotspots during the early phase of NE release (Figure 6N; Video S6), which are likely to correspond to axonal varicosities, i.e. the sites of NE release in response to neuronal activity. Analysis of the standard deviation of supervoxel responses over time revealed that these hotspots initially exhibited localized NE release and progressively diffused to a relatively uniform distribution before returning to the basal level, by comparing the data point “down to 25% of the peak” with these of “the basal” and “end” (Figures 6N and 6O).

Taken together, these results indicate that NE release is globally synchronized yet locally nuanced, suggesting that the regulatory mechanisms of the LC may operate through both brain-wide coordination and regional specificity.

### LC system globally regulates brain-wide neuronal activities in an inverted-U manner

We then investigated how the brain-wide projecting LC system affects neuronal activities across the brain. Previous studies in rodents and non-human primates showed that the LC activity level is closely linked to neural plasticity and task performance, characterized by an inverted-U curve.^1,42,44,94,95^ However, it remains unexplored whether the regulatory effects of the LC system on brain-wide neural networks are LC activity level-dependent. We thus examined brain-wide spontaneous neuronal activities across various experimental conditions corresponding to different levels of LC-NE neuronal activities: no activity (“LC ablation”), moderate level (“LC optogenetics”), and high level (“LC chemogenetics”). In comparison with the channelrhodopsin ChrimsonR-mediated optogenetic activation via 561-nm single-photon laser, TRPV1-mediated chemogenetic activation via bath application of capsaicin evoked a much higher level of long-lasting LC-NE neuronal activities and NE release (Figures S6). Three corresponding control experiments were also performed: ablation of an equal number of randomly selected non-LC cells surrounding the LC region (“Ablation control”), expression of mCherry instead of ChrimsonR in LC-NE neurons (“Opto-control”), and treatment with DMSO instead of capsaicin (“Chemo-control”). *In vivo* two-photon imaging was carried out for experiments under the LC ablation, chemogenetic activation and corresponding controls, while confocal imaging was performed for experiments under the optogenetic activation and its control.

For each experimental condition, we monitored brain-wide spontaneous neuronal Ca^2+^ activities before and after the manipulations in intact 6-dpf *Tg(elavl3:H2B-GCaMP6f)* larvae (Figure 7A), in which the calcium indicator GCaMP6f was located in the nuclear of brain-wide neurons, and examined the changes of neuronal activities in ten brain regions (Figure 7B), where the axon projections and NE release were analyzed above (see also Figures 4H, 4I and 6C). Please note, specific subregions were analyzed in some regions due to preferential locations of neuronal somata and processes in these regions, including Tel *vs* P_NL, TeO_PL *vs* TeO_NL, and H vs H_NL. Ca^2+^ activities within a 10-min window before and after LC manipulations were analyzed, with a focus on the neuronal activity strength (including the average frequency and peak amplitude of Ca^2+^ activity events, and the ratio of responsive neurons), and the correlation among all responsive neurons (Figures 7A and S7A). The modulation effects were quantified using the modulation index (MI), defined as “(After - Before)/Before”. Both LC ablation and chemogenetic activation caused a global reduction in the strength and correlation of neuronal activities across the brain, whereas LC optogenetic activation enhanced these characteristics (Figures 7C-7F).

**Figure 7.**
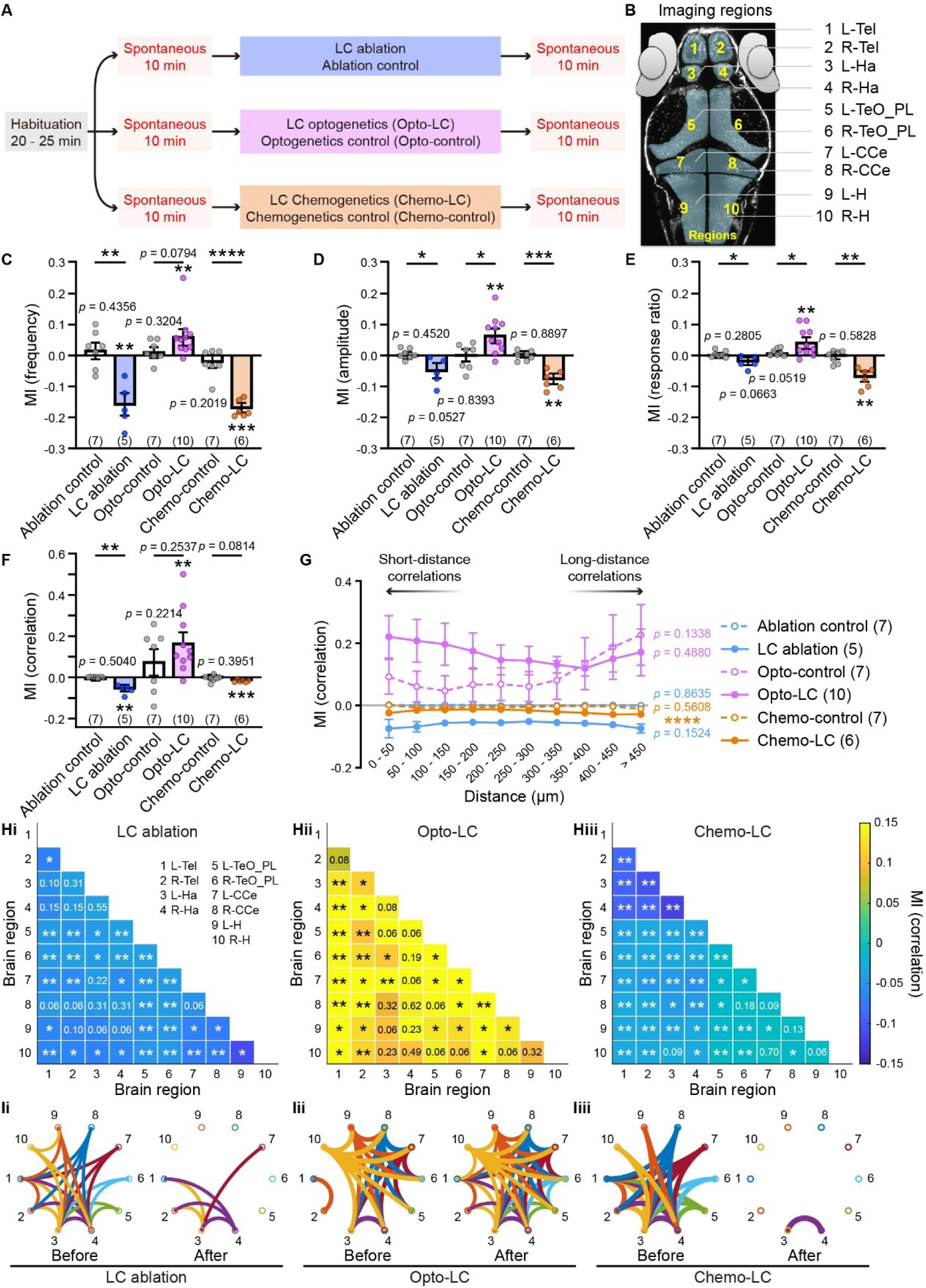
LC system globally regulates brain-wide neuronal activities in an inverted-U manner. (**A** and **B**) Overview of experimental designs and brain region segmentations for analysis of LC modulation on brain-wide neuronal activities. (**A**) Protocols for *in vivo* calcium imaging of brain-wide neuronal activities before and after LC-NE neuron manipulations. *Tg(dbh:GAL4FF,UAS:TRPV1-TagRFP);Tg(elavl3:H2B-GCaMP6f)* larvae were used for two-photon laser-based ablation and chemogenetic manipulations, and two-photon calcium imaging was performed. T*g(dbh:GAL4-VP16,UAS:ChrimsonR-tdTomato);Tg(elavl3:H2B-GCaMP6f)* larvae were used for optogenetic manipulations, and confocal imaging was performed. Larvae were habituated for 20 - 25 min prior to imaging. Spontaneous calcium activities were recorded for 10 min before and after manipulations for subsequent analysis. For control experiments, ablation of equal number of random-selected non-LC cells around LC somata, treatment with DMSO, and expression of mCherry instead of ChrimsonR in LC-NE neurons were performed, respectively. (**B**) A dorsal view of a 6-dpf *Tg(elavl3:H2B-GCaMP6f)* larva. Single-plane brain-wide calcium activities were recorded and analyzed based on brain regions, which are numbered: 1) L-Tel, left telencephalon; 2) R-Tel, right telencephalon; 3) L-Ha, left habenula; 4) R-Ha, right habenula; 5) L-TeO_PL, left optic tectum periventricular layer (i.e. TeO soma location); 6) R-TeO_PL, right optic tectum periventricular layer; 7) L-CCe, left corpus cerebelli; 8) R-CCe, right corpus cerebelli; 9) L-H, left hindbrain; 10) R-H, right hindbrain. (**C-G**) Effects of LC manipulations on brain-wide neuronal activity strength (**C-E**) and global correlation (**F**, **G**). Individual calcium activity events were detected by customed codes. For each parameter, modulation index (MI) was calculated as “(After - Before)/Before”. The number in the brackets indicate the number of larvae examined. Unpaired t-test (Welch’s t-test) was performed for comparison between corresponding experimental and control groups. Each group was also compared with 0 by one-sample t-test if it passed normality test by Kolmogorov-Smirnov test. Only the optogenetic experimental group was compared with 0 by using one-sample Wilcoxon test. (**C-F**) Modulation index of the frequency (**C**) and amplitude (**D**) of calcium events, ratio of responsive neurons (**E**), and median absolute correlation across the brain (**F**). (**G**) Modulation index of the median absolute correlation among neuron pairs at different distances. Distance bin, 50 μm. One-way ANOVA Friedman test was performed for comparison of individual experimental or control groups between different distance bins. (**H**) Modulation index of the median absolute correlations of neuron pairs from different brain regions (“inter-regional correlation”) under LC ablation (**Hi**), LC optogenetic activation (**Hii**), and LC chemogenetic activation (**Hiii**). The number on the graph represents the respective *p* value. Wilcoxon signed-rank test was performed for the optogenetic group, and Mann Whitney test was performed for the rests. (**I**) Averaged inter-regional correlations before and after LC ablation (**Ii**), LC optogenetic activation (**Iii**), and LC chemogenetic activation (**Iiii**), represented by circle graphs. Only inter-regional correlations larger than the median value of correlations before manipulations were shown. **\****p* < 0.05, **\*\****p* < 0.01, **\*\*\****p* < 0.001, **\*\*\*\****p* < 0.0001. Data are represented as mean ± SEM. See also Figures S6-S8.

The brain-wide projections of LC-NE neurons suggest their capability to modulate both local and long-range neuronal correlations. Therefore, we further analyzed neuronal correlation at varying distances or among different regions. Following three types of LC manipulations, the effects on correlations among both short-distance and long-distance neurons persistently decreased for the LC ablation and chemogenetic activation, or increased for the LC optogenetic activation (Figure 7G). Notably, the significant difference along the neuron distance for the LC chemogenetic activation was due to that the decrease of correlations among middle-distance neurons was relatively smaller than those among short- and long-distance neurons. Similar regulatory effects were observed for inter-regional neuronal correlations (Figure 7H), which were determined by calculating the correlations between neurons in paired brain regions, and the relevant functional connectivity (Figure 7I), which was depicted as circle graphs representing the averaged inter-regional correlations. For both the LC ablation control and the chemogenetic activation control, no significant changes were observed in neuronal activity strength, the correlation, or functional connectivity (Figures 7C-7G, S7B and S7D). Notably, obvious increases in the correlation but not functional connectivity were observed in the optogenetic control (Figures 7F, 7G, and S7C), which may result from the direct activation of LC-NE neurons by single-photon confocal illumination, given that these neurons exhibit light sensitivity (Figures S6C and S6F; see also Figures 2 and S2). These results together indicate that LC system regulates brain-wide neuronal activities in an inverted-U manner.

The modulatory effects of the LC depend not only on the level of NE release but also on the expression profile of NERs. Based on previously published scRNA-seq data of whole-brain cells in 8-dpf larvae,^96^ we found that both high-affinity inhibitory types (e.g., α2 receptors: adra2da) and low-affinity excitatory types (e.g., α1 receptors: adra1d; and β2 receptors: adrb2a) of NERs were predominantly expressed in inhibitory neurons rather than excitatory neurons (Figure S8A), with this pattern being comparable across various brain regions (Figure S8B). Please note that in the cerebellum, Purkinje cells are inhibitory neurons, whereas granule cells are excitatory neurons.^97^ In addition, the habenula possesses a very limited number of inhibitory neurons.^98^ Such expression profiles can well account for the observed inverted-U effects of the LC on brain-wide neuronal activities (Figure S8C; see also Figures 7C-7G). At low to moderate levels of LC activity (i.e., under spontaneous control condition or LC optogenetic activation), released NE can activate high-affinity inhibitory α2-type receptors expressed on both excitatory and inhibitory neurons. However, because inhibitory neurons express more α2-type receptors, NE at these concentrations exerts a stronger suppressive effect on inhibitory neurons, thereby enhancing the overall excitation/inhibition (E/I) ratio and neural network activity. In contrast, at moderate to high levels of LC activity (e.g., during chemogenetic activation), elevated NE release engages lower-affinity excitatory α1- and β-type receptors on both types of neuronal populations. Given the greater expression of these excitatory receptor subtypes on inhibitory neurons, this results in a stronger activation of inhibitory neurons, leading to a reduction in overall network activity. The intricate mechanisms underlying these effects need further examination.

These results together indicate that the LC globally regulates brain-wide neuronal activities in an inverted-U manner, which depends on the level of LC-NE neuronal activities.

### LC system regulates brain-wide neuronal activities with regional differences

Considering that the collective axon projection of the LC system and the release of NE exhibit bilateral symmetry but regional variations, we subsequently examined the differential regulations between the left and right hemispheres, as well as various brain regions along the anterior-posterior axis. By analyzing neuronal activities within individual brain regions, we found that intra-regional correlations in the left and right counterpart regions were equally downregulated by LC ablation and chemogenetic activation (Figures 8A and 8C, scattered groups), and equally upregulated by LC optogenetic activation (Figure 8B, scattered groups). Interestingly, the degrees of modulation varied across different brain regions (Figures 8A-8C, average groups).

**Figure 8.**
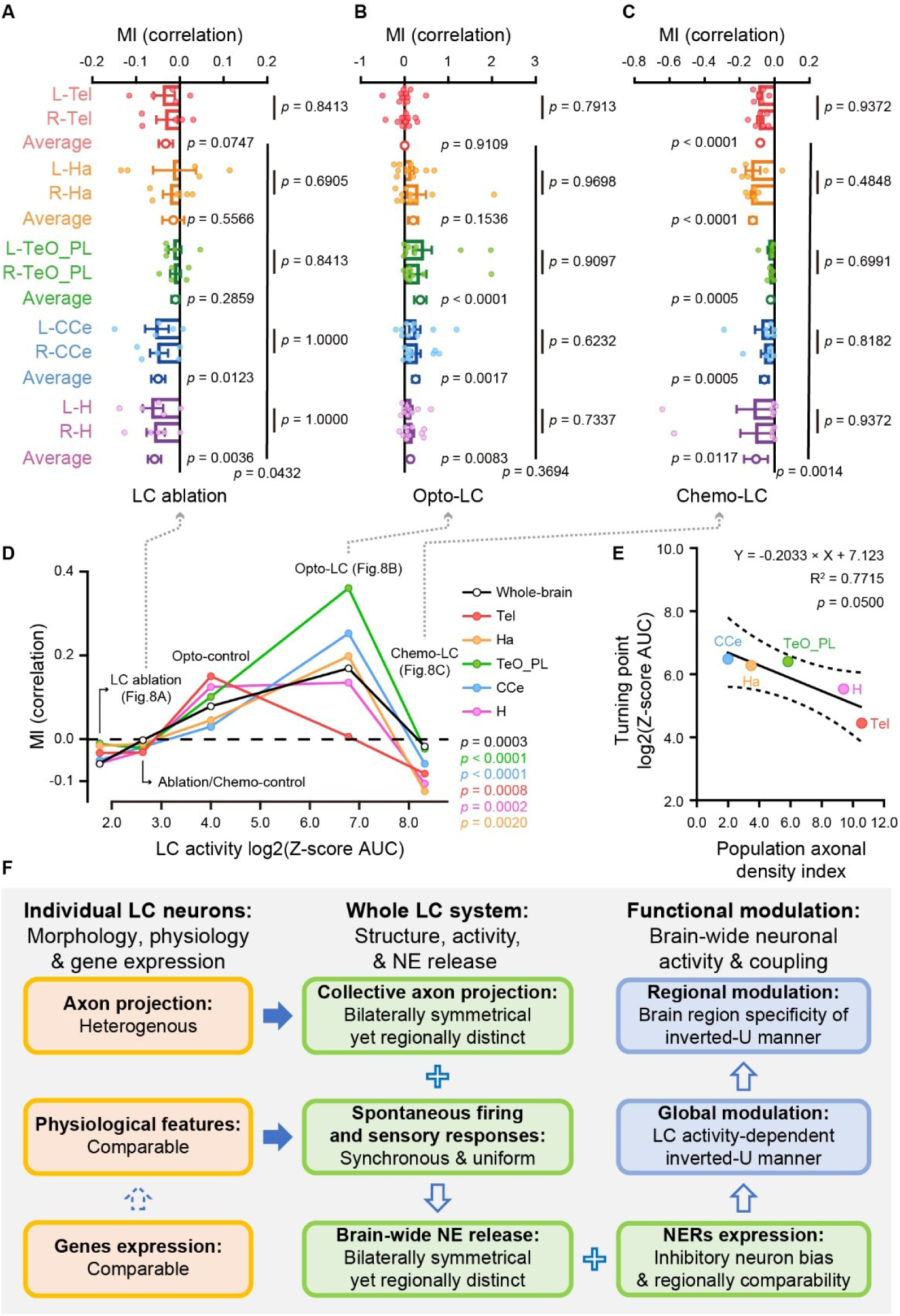
LC system modulates neuronal coupling in a region-specific inverted-U manner. (**A-C**) Modulation indices of intra-regional correlations under LC ablation (**A**), LC optogenetic activation (**B**), and LC chemogenetic activation (**C**). Each point represents the average intra-regional correlation within specific brain regions of each larva, and the average value of counterpart regions were showed as a colored circle. Wilcoxon rank sum test was used to compare bilateral counterpart regions. For comparison between the average value of counterpart regions with zero, Kolmogorov-Smirnov test was used assess normality. If the data passed this test, one-sample t-test was conducted. Otherwise, one-sample Wilcoxon signed rank test was used. (**D**) Inverted-U modulation index of intra-regional correlations under different LC activity levels induced by different experimental conditions. The LC activity, represented by Z-score AUC, was log2-transformed. Each dot represents averaged modulation index for respective regions under corresponding conditions, including LC ablation in (**A**), LC optogenetic activation in (**B**), and LC chemogenetic activation in (**C**). Ablation control and chemo-control were averaged to represent modulation index of intra-regional correlations at the basal level of LC activities. One-way ANOVA Kruskal-Wallis test was used to compare the average modulation index of each brain region under different manipulations. (**E**) Correlation between the population projection density index and the MI turning points. The data in (**D**), excluding the values in the LC ablation group, were used to estimate the turning point of the inverted-U curve. The projection density indices of population LC-NE neurons for each brain region are the same as these in Figure 4I. These turning points were fitted by Least-squares linear regression. Dotted lines show the 95% confidence bands of the best-fit line. The t-test was used to test whether the slope is significantly different from 0. (**F**) Working logic of the LC-NE system. Data are represented as mean ± SEM. See also Figures S6-S8.

To further characterize the region-specific differences of LC modulation, we plotted the modulation degree of each brain region (consisting of bilateral counterparts) against the level of LC-NE neuronal activities (see Figure S6F) under different experimental conditions. The modulation of individual regions exhibited an inverted-U manner, with variations across different regions (Figure 8D). In particular, for the telencephalon and hindbrain, the peak (“turning point”) of the inverted-U curve appeared at comparatively lower level of LC activity than other regions (Figure 8D), consistent with the results that these regions had a higher level of LC-NE neuronal axon projection’s density and NE release (see Figures 4I, 6C and 6D). Moreover, the fitted turning points of modulation degree (Figure 8D) in each region were negatively correlated with population LC-NE projection density index (Figure 8E), further supporting the presence of region-specific inverted-U modulation, with earlier turning points observed in regions with a higher LC-NE projection density. These findings suggest that the brain-wide projecting LC system can exert regionally various regulations, a characteristic that can be attributed to its regionally distinct axon projections and associated NE release.

## DISCUSSION

Our study delineates the organizational logic of the LC system by integrating three levels of examination: individual LC-NE neurons, the whole LC system, and its functional modulation (Figure 8F). First, we revealed the multidimensional characteristics of individual LC-NE neurons (Figure 8F, left). While individual LC-NE neurons exhibit heterogenous brain-wide axon projections, they share comparable physiological properties and similar expression patterns of neural function-related genes. Second, we unveiled the patterns of the structure, activity and NE release of population LC-NE neurons within the entire LC system (Figure 8F, middle). Morphologically heterogeneous LC-NE neurons assemble complementarily into the LC system, which symmetrically projects to the bilateral hemispheres with distinct targeting of brain regions along the anterior-posterior axis, and exhibits synchronized activities and uniform sensory responses, resulting in symmetrical NE release with a region-distinct pattern. Third, we demonstrated that the LC system regulates brain-wide neuronal activities in an inverted-U manner, displaying region-specific variations (Figure 8F, right). In alignment with the regionally comparable pattern of NER expression, which is biased towards inhibitory neurons, the global regulation of the LC system manifests an activity level-dependent inverted-U characteristic. This inverted-U regulation, akin to the bilaterally symmetrical yet regionally distinct pattern of population projections and NE release, displays variations among different brain regions. In conclusion, our study highlights that morphologically heterogeneous LC-NE neurons sharing comparable intrinsic properties assemble complementarily into a concerted hub to implement region-specific modulation of brain-wide neural dynamics.

Understanding of the brain-wide axon projections of individual LC-NE neurons has been limited by technical challenges.^4^ The introduction of RNA barcode technology has shed light on the projection heterogeneity of single LC-NE neurons, though it could not provide morphological structures and specific patterns in their terminal regions yet.^55^ Our study, employing genetically sparse labeling, has documented their detailed morphology and revealed morphological clusters with heterogeneous patterns of axon projections. Although technical limitations prevented from reconstructing every LC-NE neuron in individual larvae, both the total number of LC-NE neurons per larva and the relatively low proportions of certain clusters (particularly C5) suggest that not all morphological types are necessarily present in each larva. Interestingly, these morphologically heterogeneous LC-NE neurons share comparable physiological properties examined by *in vivo* whole-cell recording and comparable expression profiles of neural function-related genes examined by scRNA-seq. Thus, our study offers a multidimensional characteristic of individual LC-NE neurons.

The heterogeneous axon projections of individual LC-NE neurons provide a potential substrate for the modular organization of LC regulation observed previously.^33,58,59^ Retrograde tracing of distinct LC terminal regions often labels separate populations of LC-NE neurons, reinforcing the idea of modular organization of the LC system.^33,56,58,59^ Moreover, it was demonstrated that most LC-NE neurons in mice originate from *En1*-expressing R1 progenitor cells, with a smaller subset arising from *Hoxa2*-expressing R2 progenitor cells.^36^ Further work identified the *Pax7* marker and delineated two subpopulations of LC-NE neurons, providing additional evidence for molecular diversity.^51^ Recent transcriptomic analyses have further highlighted this heterogeneity, particularly in the expression of neuropeptides.^52-54,99^ Despite this molecular diversity, many LC-NE function-associated genes, such as *dbh*, *th* and *slc6a2*, are highly conserved across species from zebrafish to mammals.^52-54^ In combination with the conserved functions of the LC system, our findings from zebrafish may have broad implications and provide insights that are translatable to the LC system in other vertebrates.

In addition to its structural projection pattern, the output of the LC system is shaped by the activity patterns of LC-NE neurons. We found that population LC-NE neurons show synchronized spontaneous activities and uniform sensory responses. Consistent with previous studies showing that gap junctions of LC-NE neurons facilitate their synchronous firing and response to salient stimuli,^23,35,91^ we found that the gap junctions exist among LC-NE neurons of zebrafish, especially in ipsilateral LC-NE neurons (Figure S5). As the somata of LC-NE neurons are segregated, the gap junctions probably exist in local dendrites among ipsilateral LC-NE neurons. The concentrated dendrites around the soma may receive common synaptic inputs as well as electrical coupling via gap junctions, contributing to the firing synchrony of ipsilateral LC-NE neurons.^34,35^ As the ratio of electrical coupling between bilateral LC-NE neurons is quite low, we speculate that the firing synchrony of bilateral LC-NE neurons may be largely attributable to common synaptic inputs, a notion consistent with the finding of convergent inputs onto LC-NE neurons.^10^

Supporting the well-documented inverted-U relationship between LC activity and behavioral performance,^1,42,94^ our study demonstrated that the regulatory effects of the LC system on whole-brain neuronal activities are also dependent on the level of LC-NE neuronal activities, offering the first evidence for the neuronal basis of the inverted-U regulation. The effect of NE released by LC-NE neurons depends on the types of NERs expressed on downstream neurons. It is known that α2-type NERs (including α2A, α2B, α2C and α2D) have a higher affinity for NE than α1-type NERs (including α1A, α1B and α1D), and both of these NERs have higher affinities for NE than β-type NERs (including β1, β2 and β3).^95,100,101^ Furthermore, the activation of α2 receptors inhibits neuronal activities, whereas α1 and β receptors are excitatory.^95,101^ Based on the inverted-U regulation of whole-brain neuronal activities, we speculate that NERs may preferentially express on inhibitory neurons. This would lead to increased activities of whole-brain neural networks when low levels of NE activate α2 receptors on inhibitory neurons. Conversely, high levels of NE, associated with high-level LC activities, would activate α1 and/or β receptors on inhibitory neurons, suppressing neural network activities. As expected, our scRNA-seq analysis revealed that inhibitory neurons indeed exhibit higher expression of α2, α1 and β receptors than excitatory neurons across the brain (see Figure S8). Future studies are needed to investigate the intricate mechanisms underlying these effects.

Early studies posited that the LC functions as a relatively homogeneous nucleus in regulating diverse neural functions and behaviors.^1,2,49,50^ However, technological advancements have unveiled a more complex picture, highlighting the heterogeneous nature of the LC and its capacity for modular regulation.^4,13,32,33,56,58-61^ This growing body of evidence, initially inferred from molecular diversity,^51-54^ variability in LC-NE neuronal activity patterns,^59-61,102^ and axon projection-specific functions,^33,58,59,64^ collectively challenges the notion of LC homogeneity and supports the concept of a modular, heterogeneous system.

Our study contributes to this evolving understanding by offering a comprehensive, multidimensional characterization of the LC system. We uncovered both homogeneous and heterogeneous aspects of the LC system. On one hand, our findings shed light on the homogeneous aspects of the LC system, including the brain-wide axon projections, comparable physiological properties and relevant genes’ expression of individual LC-NE neurons, the symmetry of collective axon projections, the firing synchrony of population LC-NE neurons, synchronized global NE release, and the brain-wide regulation of neuronal activities. On the other hand, we provide a detailed analysis of individual LC-NE neuron heterogeneity, particularly their diverse axon projection preferences, which likely form the structural basis for modular regulation. Moreover, we demonstrated the region variations of collective axon projections, NE release, and functional regulation. This region-specific heterogeneity likely contributes to the modular regulation of the LC system. Thus, our findings offer a systematical perspective on LC organization, emphasizing the crucial role of region-specific heterogeneity in understanding LC modularity and its functional implications.

Our findings suggest a potential dual mechanism by which the LC regulates brain functions. From the perspective of homogeneity, the LC broadcasts signals to its widespread projection targets, simultaneously influencing network activities across the brain. Such global upregulation or downregulation, determined by the level of the LC system, follows an inverted-U pattern across the brain (see Figure 7). In contrast, from the perspective of heterogeneity, the LC system displays region-specific modulation (see Figures 8A-8E), characterized by spatial variability in different brain regions. Our work thus provides multifaceted empirical support and an integrated framework for understanding the dual characteristics of the LC—its capacity for both global homogeneity and local or regional heterogeneity. These findings lay a foundation for future investigations into context-dependent, region-specific LC modularity during specific cognitive processes.^13^

### Limitations of the study

While our study provides a comprehensive examination of certain aspects of LC heterogeneity, its full scope likely extends beyond the parameters we investigated. Several factors, in combination with the identified heterogeneity, could further expand the diverse functional roles of the LC system. First, variations in the density of LC axon varicosities, along with differences in the types and distributions of NERs on different brain cells (e.g., neurons, glia, as well as vascular cells), interact with localized modulation in specific brain regions. This interplay could significantly amplify the functional diversity of the LC within specific regions or neural circuitries.^4,17,24,45,69,79,95,103-105^ Second, the potential shift from synchronous to asynchronous discharge patterns in adult mammals may further contribute to the functional heterogeneity of LC outputs.^9,61,62,91,102^ Third, although the LC system exhibits broad projections and NE release, the co-release of other neurotransmitters, such as galanin, dopamine, or glutamate, could enrich distinct regulatory functions.^52,53,68,106^ Fourth, LC heterogeneity appears to increase along evolution, likely in parallel with the increasing complexity of cognitive functions in animals. In large-brained species such as primates, the absolute number of LC-NE neurons increases, whereas their proportion relative to total neurons in the brain decreases.^4,107^ This disparity implies a shift toward more nuanced and spatially specific modulation of the LC system through increased diversity in molecular expression, axonal projections as well as neuronal activity patterns.^12,33,63^ All these suggest that LC heterogeneity may operate at multiple levels, explaining how the LC system can differentially regulate a wide array of brain functions.

## RESOURCE AVAILABILITY

### Lead contact

Further information and requests for resources and reagents should be directed to and will be fulfilled by the lead contact, Jiu-Lin Du.

### Materials availability

Plasmids and fish lines generated in this study are available from the lead contact without restriction.

### Data and code availability

- The raw sequence data reported in this paper have been deposited in the Genome Sequence Archive in National Genomics Data Center, China National Center for Bioinformation/Beijing Institute of Genomics, Chinese Academy of Sciences (GSA: CRA023422) that are publicly accessible at https://ngdc.cncb.ac.cn/gsa.
- This paper does not report any original code.
- Any additional information required to reanalyze the data reported in this work paper is available from the lead contact upon request.

## ACKNOWLEDGMENTS

We are grateful to Dr. Herwig Baier for providing the *Tg(UAS:Kaede)*, Dr. Misha Ahrens for providing the *Tg(elavl3:H2B-GCaMP6f)*, Dr. Koichi Kawakami for providing *Tg(UAS:GCaMP6s)*, Dr. Marnie Halpern for providing *Tg(4×nrUAS:GFP)*, Dr. Qian Hu for optical imaging support, Dr. Zhiyong Liu, Dr. Chao Li and Dr. Min Zhang and the platform for single-cell sequencing support, Dr. Chenyang Li, Dr. Yiheng Zhang, Dr. Qiusui Deng, Dr. Sha Li, Dr. Qiyuan Zhuang, Dr. Jiajia Ye, Yu Qian, Yun Du, and Yingling Chen for their help on the morphological reconstruction of LC-NE neurons, Minjia Chen for constructing the plasmids *dbh:GAL4-VP16,UAS:ChrimsonR-tdTomato* and *dbh:GAL4FF,UAS:TRPV1-TagRFP*, Dr. Yuxi Li for sharing the NBLAST codes, Dr. Peishan Xiang and Dr. Qiusui Deng for their help on the LC tissue dissection, Dr. Yufan Wang for her help on light-sheet imaging, Hao Zhai (Institute of Automation, CAS) for his help on the video denoising using an open-source Real-ESRGAN-General-x4v3 model. This work was supported by National Science and Technology Innovation 2030 Major Program of the Ministry of Science and Technology (2021ZD0204500 and 2021ZD0204502), Science Fund for Creative Research Groups (32321003) and Key Joint Funds for Cooperation (U22B200571) of the National Natural Science Foundation of China, Shanghai Municipal Science and Technology Major Project (18JC1410100, 2018SHZDZX05 and 2019SHZDZX02), and Key Research Program of Frontier Sciences (QYZDYSSW-SMC028) and Strategic Priority Research Program (XDB32010200) of Chinese Academy of Sciences.

## AUTHOR CONTRIBUTIONS

J.D. and C.Z. conceived and supervised the project, and designed the experiments. C.Z. performed most of the experiments and analyzed data. Y.Y. performed electrophysiological experiments, calibration of reconstructed LC-NE neuronal morphology and immunohistochemistry. F.L. analyzed the data of brain-wide neuronal activities and generated the *Tg(elavl3:NES-jRGECO1a)* line. H.Z. analyzed the scRNA-seq data. L.S. examined MO-NE neuronal projections. J.L. generated the *Ki(dbh:GAL4-VP16)* F0 founder animals, and C.Z performed founder screening to identify germline-transmitting lines and generated the *Tg(dbh:GAL4-VP16,UAS:ChrimsonR-tdTomato)* and *Tg(dbh:GAL4FF,UAS:TRPV1-TagRFP)* lines. X.D. generated the *Tg(UAS:sypb-EGFP)* line. C.Y. and Z.Y. helped deploy the image registration algorithm. J.F. and Y.L. provided GRAB sensors. J.H., R.Z. and X.D. participated in discussion. J.D. and C.Z. wrote the paper with inputs from F.L., Y.Y. and H.Z.

## DECLARATION OF INTERESTS

The authors declare no competing interests.

## SUPPLEMENTAL FIGURE LEGENDS

**Figure S1.**
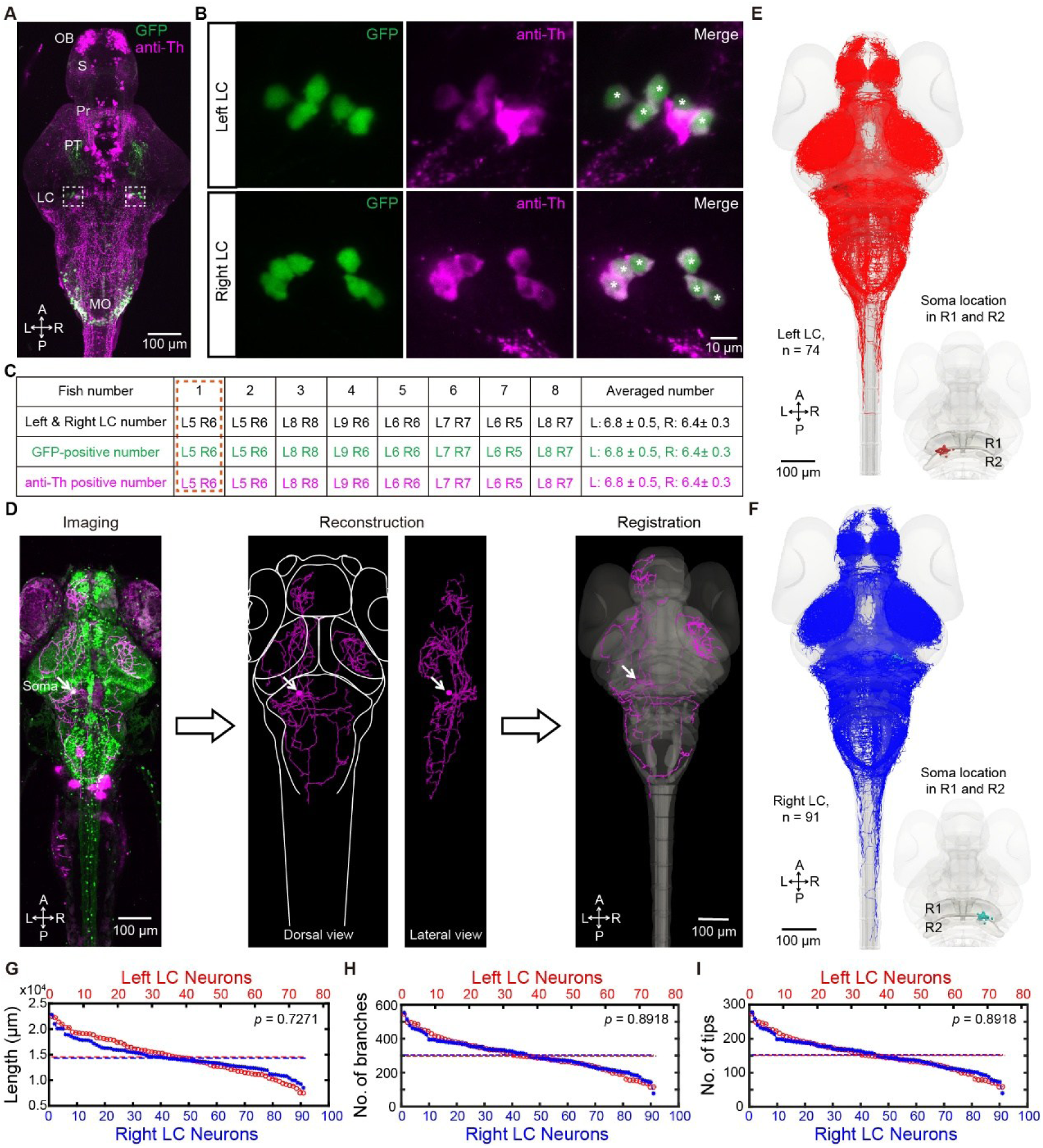
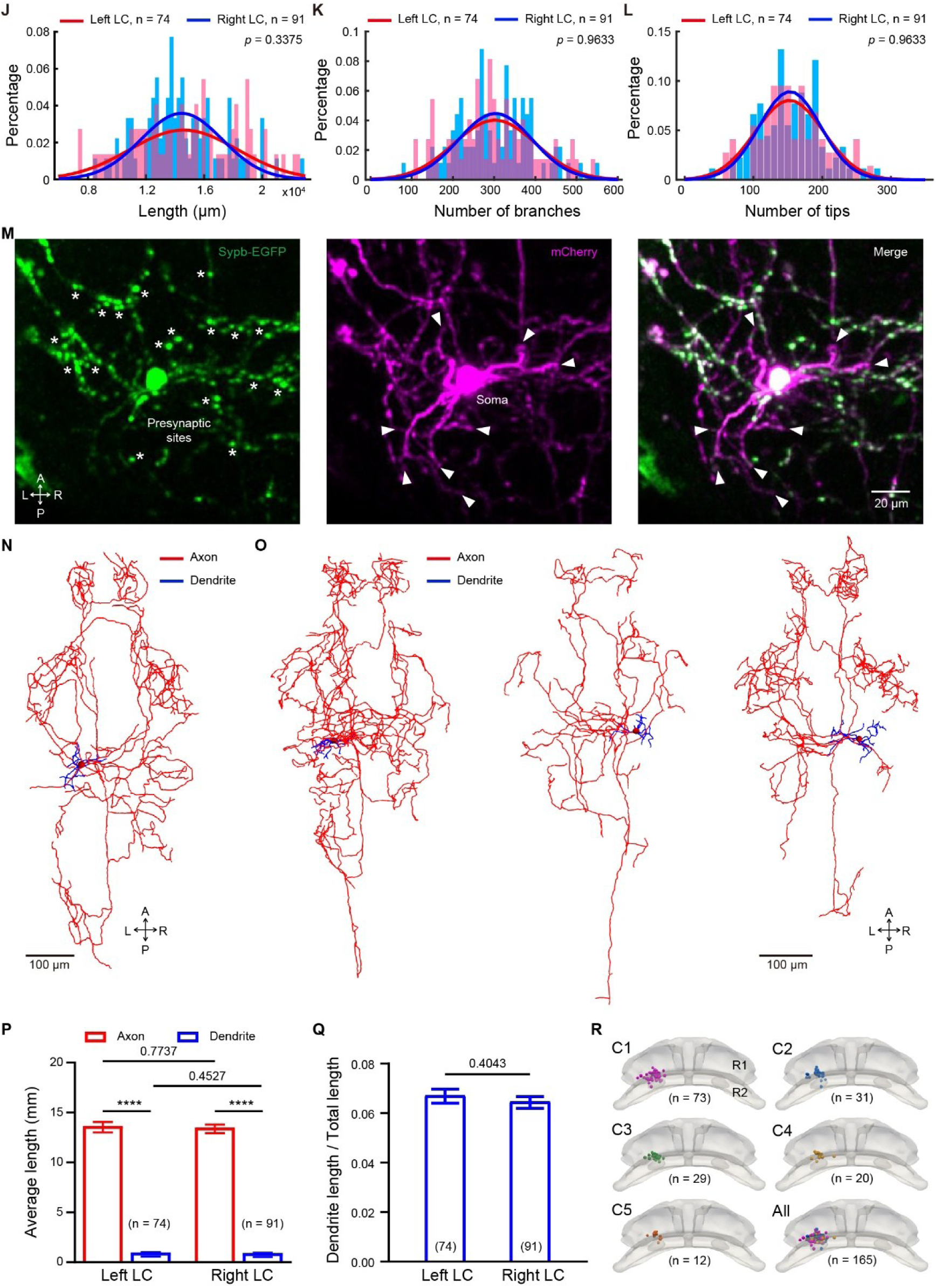
Reconstruction and characterization of individual LC-NE neurons’ morphology reveal that bilateral LC-NE neurons exhibit similar complexity, related to Figure 1. (**A**-**C**) Immunohistochemical verification of the identity and total number of LC-NE neurons in 6-dpf *Ki(dbh:GAL4-VP16);Tg(4×nrUAS:GFP)* larvae by immunostaining of tyrosine hydroxylase (Th). Th-positive neurons (dopaminergic neurons and NE neurons) are in magenta, whereas dbh-driven GFP expression specifically labels NE neurons in green. (**A**) A dorsal view of 3D projection of a 280-μm stack. OB, olfactory bulb; SP, subpallium; PreT, pretectum; PT, posterior tuberculum; LC, locus coeruleus; MO, medulla oblongata. A, anterior; P, posterior; L, left; R, right. (**B**) A zoomed-in view of the soma regions of the left (top) and right LC (bottom), indicated by the dashed squares in (**A**), showing 100% co-localization of Th-positive cells and GFP-positive cells in the LC. Notably, Th-positive signal was predominantly localized to the cytoplasm, while GFP signal was distributed in both cytoplasmic and nuclear compartments. Asterisks mark LC-NE somata. (**C**) Summary of the number of GFP-positive and Th-positive neurons in unilateral LC. Left LC, 6.8 ± 0.5; right LC, 6.4 ± 0.3. The data were obtained from 8 larvae at 6 dpf. The data shown in (B) are from the larva 1. (**D**-**F**) Reconstruction of 165 individual LC-NE neurons, each of which was obtained from an individual *Ki(dbh:GAL4-VP16);Tg(UAS:Kaede);Tg(elavl3:GCaMP5G);Et(vmat2:EGFP)* larva at 6 dpf. (**D**) Pipeline of the morphological reconstruction of individual LC-NE neurons. Left, dorsal view of 3D projection of a typical single LC-NE neuron (magenta) with reference signals (green). Middle, 3D reconstruction of the LC-NE neuron’s morphology based on Kaede fluorescent signal (magenta). Right, registration of the reconstructed morphology of the LC-NE neuron into the brain template based on green reference signals. White arrows indicate the soma of the reconstructed LC-NE neuron. (**E** and **F**) Projections of left 74 LC-NE neurons (**E**) and right 91 LC-NE neurons (**F**) in the brain template. The bottom right images indicate the location of LC-NE neuronal somata. (**G-I**) Characterization of the morphological features of the individual LC-NE neurons, including the total length (**G**), number of branches (**H**), and number of branch tips (**I**), shown for 74 left (red) and 91 right (blue) LC-NE neurons in a descending order. The dashed lines indicate the respective mean values. (**J-L**) Normal distribution of projection features for left (light red) and right (light blue) LC-NE neurons, including the total length (**J**), number of branches (**K**), and number of branch tips (**L**). Normal distributions were fitted using MATLAB’s fitdist function with maximum likelihood estimation (MLE). The *p* value indicates no significant difference between the left and right LC-NE neurons. (**M-Q**) Discrimination of axons and dendrites of LC-NE neurons. (**M** and **N**) Simultaneous genetic labeling of presynaptic sites and intact morphology of a single LC-NE neuron. Data were obtained from a 6-dpf *Ki(dbh:GAL4-VP16);Tg(UAS:mCherry);Tg(UAS:sypb-EGFP)* larva. (**M**) *In vivo* confocal imaging of the morphology (mCherry signal, magenta) and presynaptic sites (EGFP signal, green) of a single LC-NE neuron. Left, EGFP-labeled presynaptic sites on axons. Middle, mCherry-labeled morphology including axons and dendrites. Right, merged signals. The asterisks indicate some of the varicosities on LC-NE neuronal axons, and the arrows mark LC-NE neuronal dendrites branches. The full morphological reconstruction of this neuron is shown in (**N**). Please note, the dendrite also expresses relatively weak EGFP due to overexpression effects. The axon is easily identified based on the full morphology reconstruction, and the dendrite is discriminated by multiple features, including no or week EGFP expression and short branch length. (**O**) Empirically discriminated dendrites and axons of individual LC-NE neurons according to the dendrites features, with dendrites colored in blue and axons colored in red. Three examples of one left LC-NE neurons and two right LC-NE neurons were shown. (**P**) Average lengths of axons and dendrites per cell of 165 reconstructed LC-NE neurons. Data were obtained from 74 left and 91 right LC-NE neurons. Paired t-test for unilateral LC-NE neurons and unpaired t-test for bilateral LC-NE neurons in length comparison. (**Q**) Average ratio of dendrite length to total length of LC-NE neuronal processes. (**R**) Somatic distribution of each morphologically defined cluster of LC-NE neurons in the rhombencephalon 1 (R1) and rhombencephalon 2 (R2) regions. The somata from all clusters were merged in the final image. C1 - C5, cluster 1 to cluster 5. The *p* values were determined using paired t-test (**C**), unpaired Wilcoxon rank sum test (**G-I**), unpaired t-test (**Q**). The data follows a normal distribution tested by Kolmogorov-Smirnov test (**J-L**), and two-sample Kolmogorov-Smirnov test for the distribution comparison (**J-L**). **\*\*\*\****p* < 0.0001. Data are represented as mean ± SEM.

**Figure S2.**
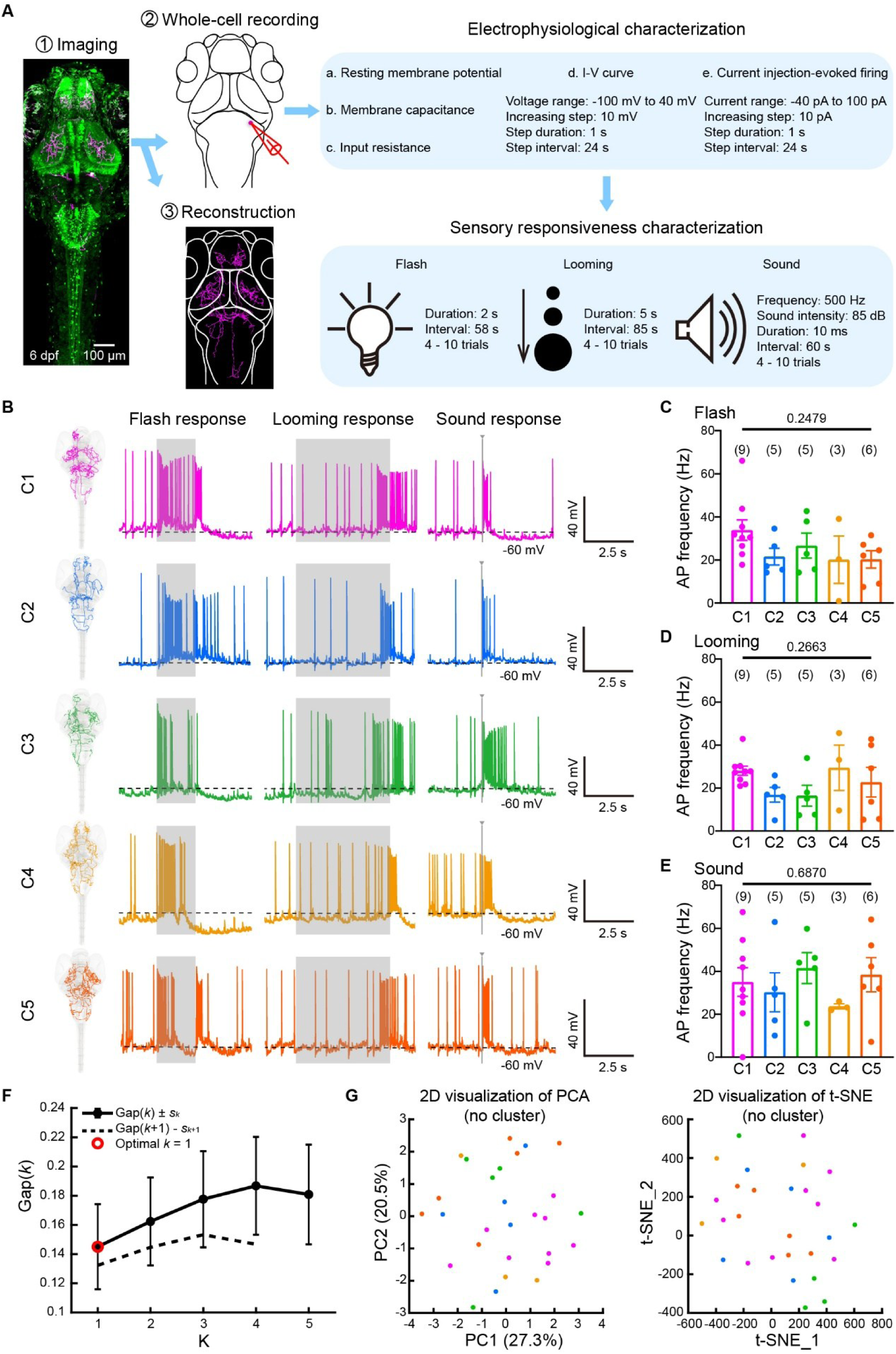
Protocol for whole-cell recording, characterization of sensory responses, and gap statistic analysis of physiological properties of LC-NE neurons, related to Figure 2. (**A**) Schematic of whole-cell recording of LC-NE neurons after morphology imaging on *Ki(dbh:GAL4-VP16);Tg(UAS:Kaede);Tg(elavl3:GCaMP5G);Et(vmat2:EGFP)* larvae. The morphological data of LC-NE neurons obtained by *in vivo* confocal imaging (process 1) at 6 dpf were later reconstructed (process 3) for subsequent morphological analysis. After the morphological imaging, part of neurons were randomly selected for *in vivo* whole-cell recording at 6 - 8 pdf (process 2) for sequentially examining intrinsic electrophysiological properties and sensory responses. The intrinsic electrophysiological properties were examined via the voltage clamp (from -100 mV to 40 mV with 10-mV step lasting 1 s) and current clamp (from -40 pA to 100 pA with 10-pA step lasting 1 s), and sensory responses were evoked by 2-s flash, 5-s looming and 10-ms sound stimuli. (**B-E**) Sensory stimulus-evoked firing activities of individual LC-NE neurons of C1 - C5. Data were obtained from 28 LC-NE neurons. (**B**) Representative cases of whole-cell recordings from C1 - C5. Grey shadows indicate the time window of sensory stimuli (2-s flash, 5-s looming, and 10-ms pure tone). (**C-E**) AP frequency of responses evoked by flash (**C**), looming (**D**), and pure tone (**E**) in LC-NE neurons of C1 - C5. Data were obtained from 28 LC-NE neurons. Each data point represents the average value of 4 - 10 trials for each neuron. The number in the brackets represents the number of neuron examined. One-way ANOVA test was performed. (**F**) Estimation of the optimal number of clusters. Gap statistic analysis was used to estimate the optimal number of clusters by comparing the log within-cluster dispersion of the observed data to that of a reference null distribution, averaged over multiple uniform resampling replicates.^89^ The optimal *k* was defined as the smallest *k* such that Gap(*k*) ≥ Gap(*k*+1) − *s_k_*_+1_. The optimal number of cluster is 1 for the 28 LC-NE neurons examined, each characterized by 5 electrophysiological properties and 6 sensory response properties (see Methods). K, number of clusters. (**G**) Two dimensionality reduction methods, principal component analysis (PCA, left) and t-distributed stochastic neighbor embedding (t-SNE, right), were applied to visualize the property profiles of the 28 LC-NE neurons characterized with 11 parameters. Each point represents the data obtained from an individual neuron. Data are represented as mean ± SEM.

**Figure S3.**
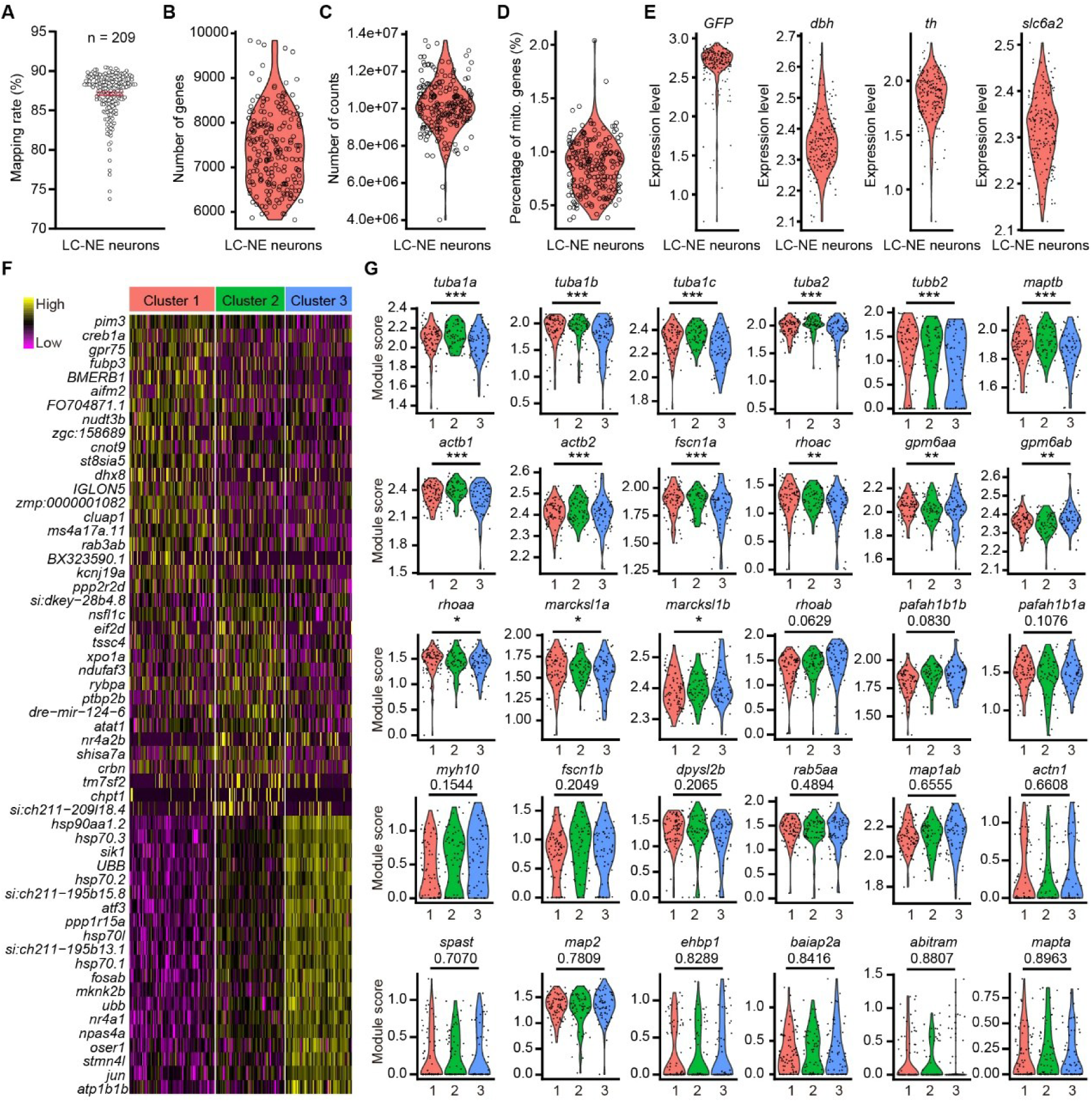
Gene expressions of individual LC-NE neurons show differences in development-related genes, related to Figure 3. (**A**) Mapping rate of 209 LC-NE neurons. Average mapping rate is 87.1%. (**B-D**) Quality assessment of the single-cell sequencing data of individual LC-NE neurons. (**B**) Number of unique genes detected within individual LC-NE neurons. (**C**) Number of genes detected within the individual LC-NE neurons. (**D**) Percentage of reads that map to the mitochondrial genome of individual LC-NE neurons. (**E**) Expression level of typical marker genes of LC-NE neurons. (**F**) Heatmap of top 20 marker genes among 3 clusters. (**G**) Expression of cytoskeletal organization-related genes among 3 clusters. One-way ANOVA test was performed (**G**). **\****p* < 0.05, **\*\****p* < 0.01, **\*\*\****p* < 0.001.

**Figure S4.**
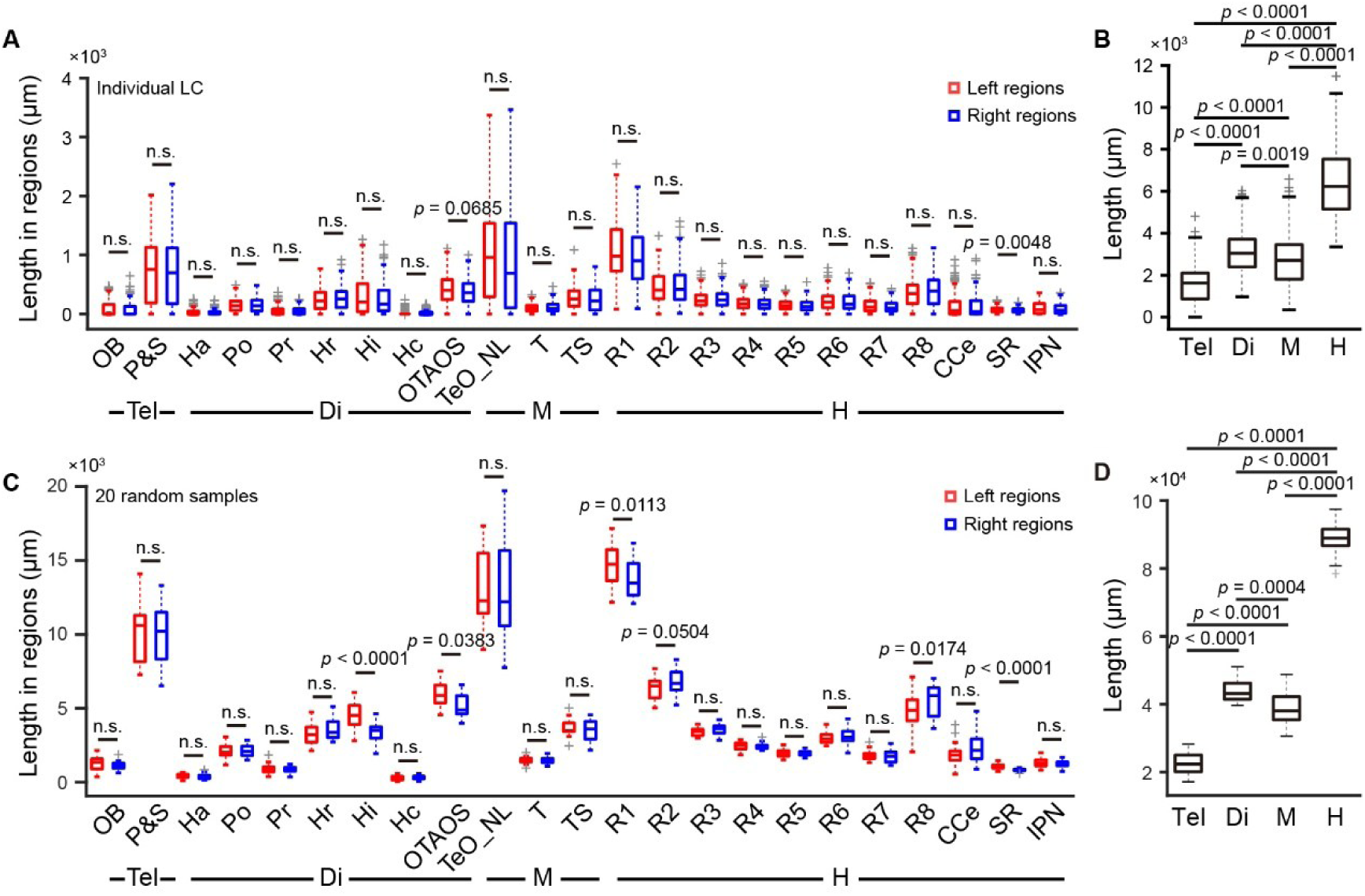
Projection length distribution of population LC-NE neurons in each brain regions of bilateral hemispheres, related to Figure 4. (**A** and **B**) Projection distribution of collective LC-NE neurons reconstructed. The dataset is the same as that in Figures 4C and 4D. (**A**) Average length of LC-NE neuronal projections in each brain region of the left (red) and right (blue) hemispheres. This was calculated as the average length of all 74 left LC-NE neurons and 74 randomly selected neurons from 91 right LC-NE neurons. The median was represented by a central line, while the interquartile range (IQR) was defined by the first and third quartiles, displayed as box edges. The whiskers were extended to the most extreme non-outlier data points, and values exceeding this range were marked as outliers (grey markers). Tel, telencephalon; Di, diencephalon; M, mesencephalon; H, hindbrain. (**B**) Average projection length of all 165 LC-NE neurons in brain areas along the anterior-posterior axis. (**C** and **D**) Simulation of the entire LC system in an individual larva by cluster-based proportional random sampling from 165 individual LC-NE neurons reconstructed. (**C**) Projection length of the simulated entire LC system in each brain region of the left (red) and right (blue) hemispheres. (**D**) Projection length of the simulated entire LC system in brain areas along the anterior-posterior axis. For two-group comparisons, a paired t-test was used for normally distributed data, while the Wilcoxon matched-pairs signed-rank test was applied for non-normally distributed data (**A**, **C**). Unpaired t-test was applied for inter-regional comparisons (**B, D**). n.s., not significant.

**Figure S5.**
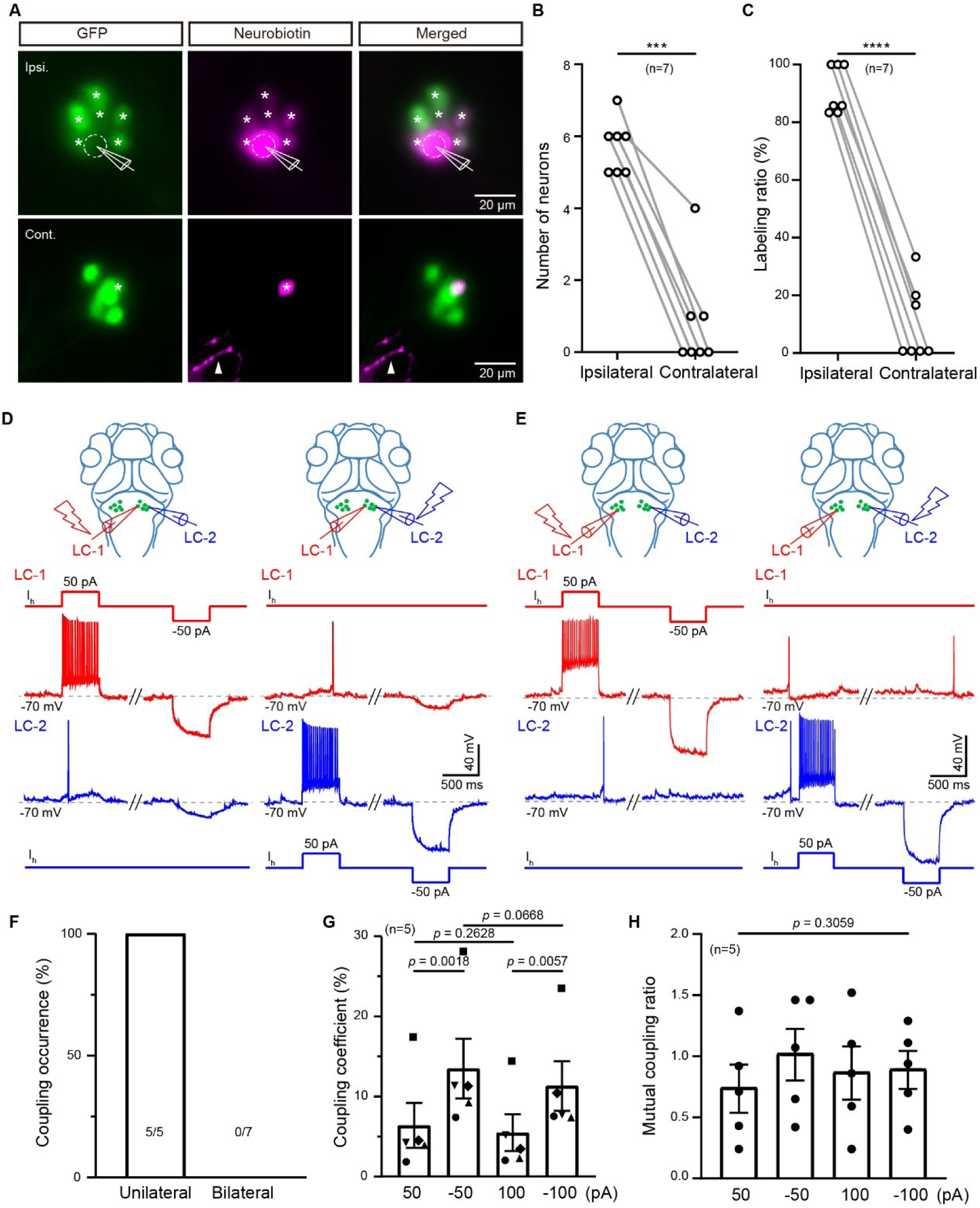
Unilateral LC-NE neurons exhibit electrical coupling, related to Figure 5. (**A-C**) Gap junctions among LC-NE neurons, revealed by neurobiotin-loading of a single LC-NE neuron in *Ki(dbh:GAL4-VP16);Tg(4×nrUAS:GFP)* larvae at 6 dpf. (**A**) Representative of neurobiotin-loading experiments. Neurobiotin (2.5%) in intracellular solution was loaded into one left LC-NE neuron (top, white dashed circle) via whole-cell recording micropipette and visualized by fluorescent immunostaining (magenta). Neurobiotin signals were observed in all 6 left LC-NE neurons (top, asterisks) and one of 5 right LC-NE neurons (bottom, asterisks). The arrowhead indicates the neurites of the recorded left LC-NE neuron. Weak GFP fluorescent signals in the recorded neuron might be due to neurobiotin-loading manipulation. (**B** and **C**) The number (**B**) and percentage (**C**) of neurobiotin-labeled ipsilateral and contralateral LC-NE neurons after one LC-NE neuron was loaded with neurobiotin. Data were collected from 7 larvae. (**D-H**) Electrical coupling among LC-NE neurons, revealed by dual whole-cell recordings of unilateral or bilateral pairs of LC-NE neurons in *Ki(dbh:GAL4-VP16);Tg(4×nrUAS:GFP)* larvae at 6 dpf. (**D** and **E**) Representative of dual whole-cell recordings of a unilateral (**D**) or bilateral (E) pair of LC-NE neurons. In current clamp mode (I = 0 pA), 0.5-s current steps at ± 50 pA currents were injected into one recorded neuron (lightning symbol), while depolarization or hyperpolarization was recorded in both neurons, and vice versa. Top: schematic of dual recordings. Bottom: current injection protocols and membrane potential traces of the pair of recorded neurons. (**F**) Coupling occurrence of unilateral or bilateral pairs of recorded LC-NE neurons. Data were recorded from 5 unilateral and 7 bilateral pairs. (**G**) Coupling coefficency of unilateral pairs of recorded LC-NE neurons, calculated as the ratio of voltage changes between un-injected and injected neurons with step currents of ± 50 pA and ± 100 pA. Each point represents averaged coupling coefficency of a pair of recorded neurons. Data were obtained from 5 pairs of unilateral LC-NE neurons. (**H**) Mutal coupling ratio of unilateral pairs of recorded LC-NE neurons, calculated as the ratio of coupling coefficency of a pair of recorded neurons with step currents of ± 50 pA and ± 100 pA. The dataset is the same as that in (**G**). Paired t-test was performed (**B**, **C**, **G**). One-way ANOVA test was performed (**H**). \*\*\*\**p* < 0.0001. Data are represented as mean ± SEM.

**Figure S6.**
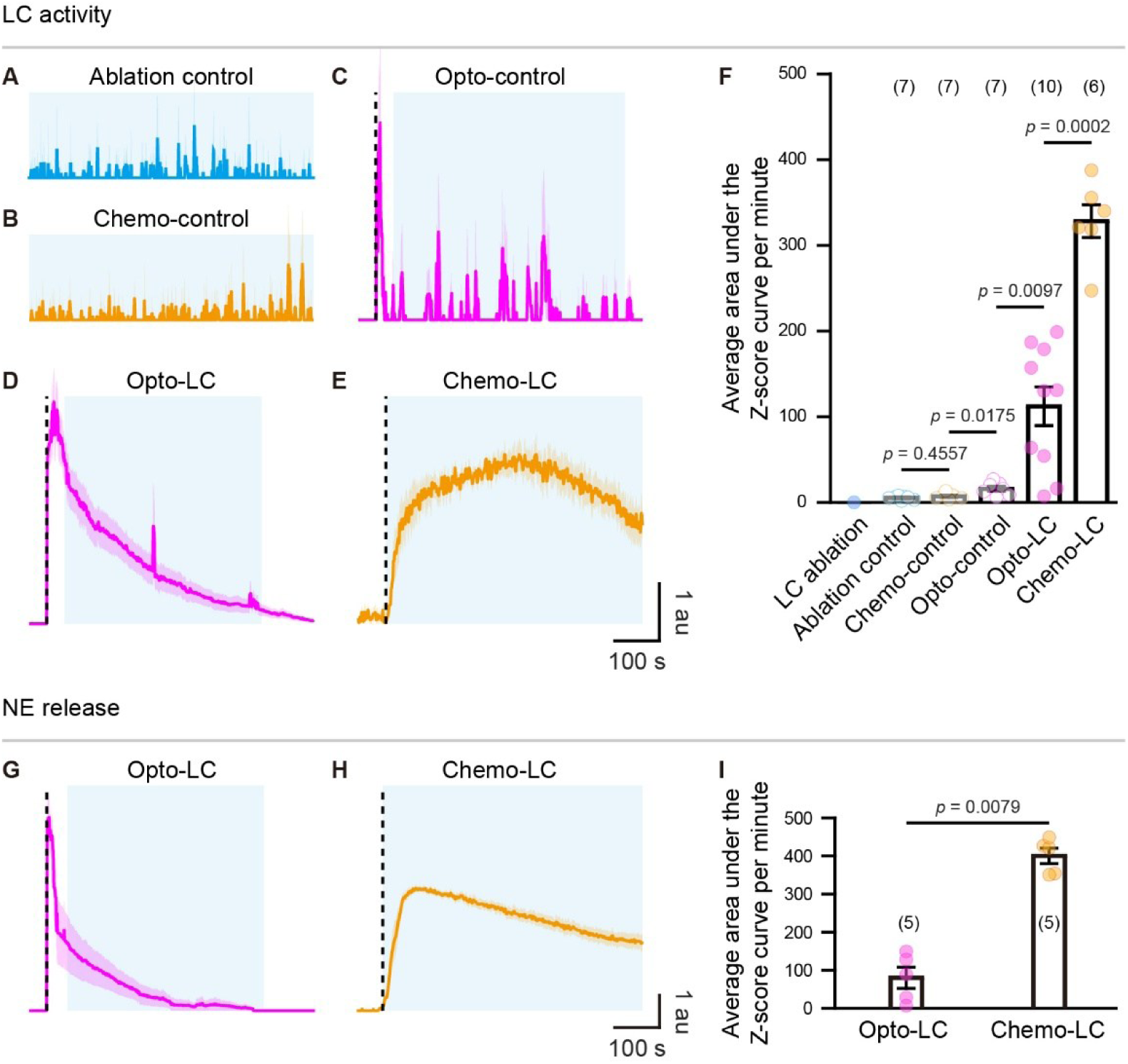
LC activities and NE release levels under different experimental conditions, related to Figures 7 and 8. (**A-F**) Averaged bilateral LC activity under different LC manipulations. (**A-E**) Averaged baseline-corrected Z-score traces of bilateral LC activities under the ablation control (**A**), chemo-control (**B**), opto-control (**C**), opto-LC (**D**), and chemo-LC (**E**). The light blue shadow indicates the time window used for calculating the area under the curve (AUC), the same window used for brain-wide neuronal activity analysis in Figures 7, S7 and 8. Dashed lines mark the optogenetic stimulation onset in (**C**, **D**) and the response onset in (**E**). Please note, the light response of LC-NE neurons evoked immediately by single-photon optogenetic stimuli was excluded from analysis in (**C**, **D**). First, the change in fluorescence (ΔF/F_0_) was calculated, and significant responses were detected using custom programs. The ΔF/F_0_ values were then transformed into Z-scores to facilitate normalization and comparison of different functional profiles derived from confocal or two-photon imaging. The baseline of the Z-score traces was corrected to eliminate negative values. The area under the baseline-corrected Z-score curve (Z-score AUC) was calculated, with only the AUC of significant responses contributing to the final result for statistical analysis. (**F**) Summary of data. Each point represents the averaged Z-score AUC of individual larvae. The number in the brackets indicates the number of larvae examined. A Mann-Whitney test was used for statistical analysis. (**G-I**) Averaged NE release within unilateral TeO_NL under different LC manipulations. (**G** and **H**) Averaged baseline-corrected Z-score traces of NE release under the opto-LC (G) and chemo-LC (**H**). The light blue shadow indicates the time window used for calculating the Z-score AUC. Dashed lines mark the optogenetic stimulation onset in (**G**) and the response onset in (**H**). Similar to (**C**, **D**), the component of NE release immediately following the single-photon optogenetic stimulus was excluded from analysis in (**G**), since this component can be due to direct light responses of LC-NE neurons evoked by the optogenetic stimuli. (**I**) Summary of data. Each point represents the averaged Z-score AUC of individual larvae. The number in the brackets indicates the number of larvae examined. A Mann-Whitney test was used for statistical analysis. Data are presented as mean ± SEM.

**Figure S7.**
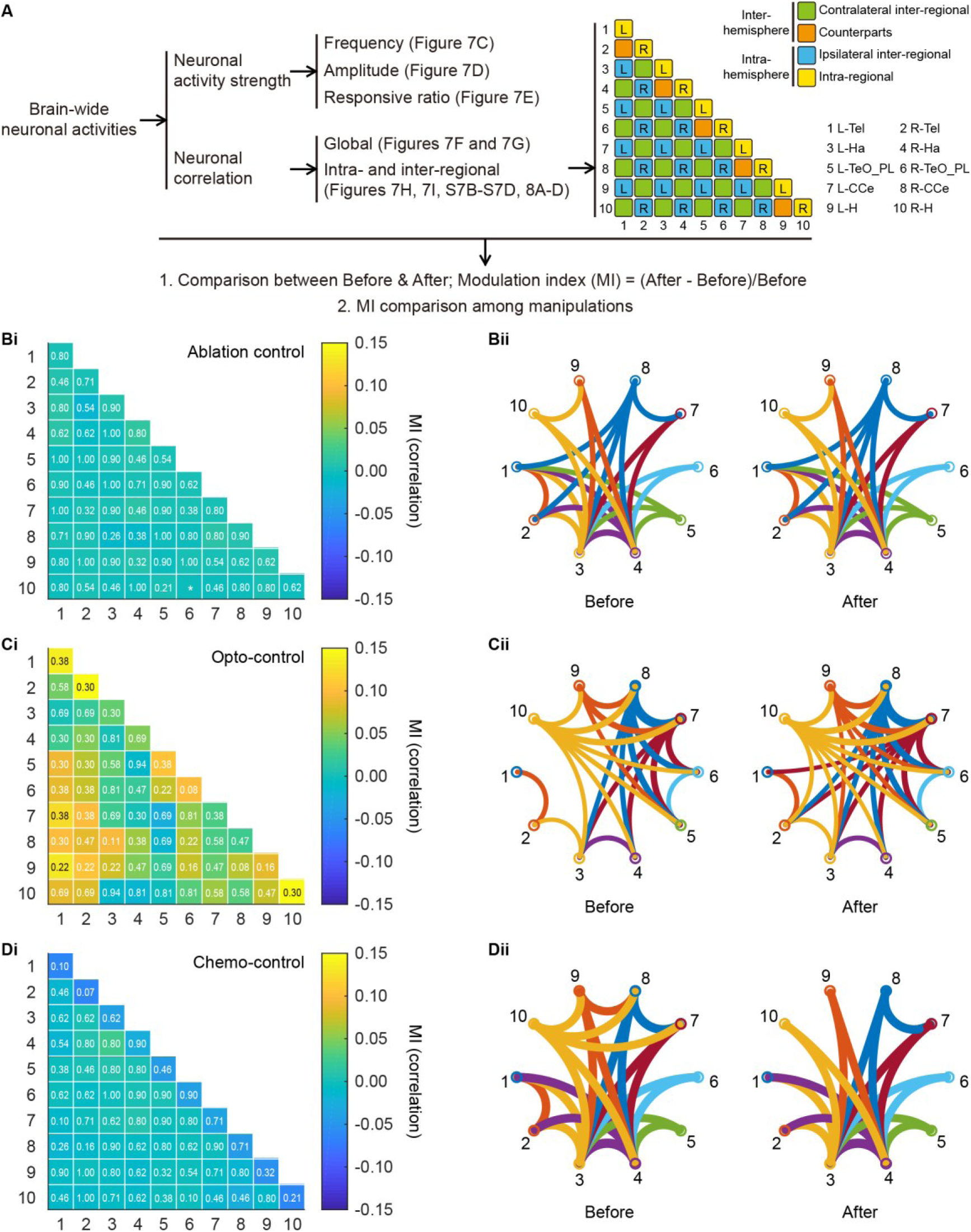
Control experiments of LC manipulations do not significantly affect network coupling, related to Figures 7 and 8. (**A**) Schematic workflow for analyzing calcium activities of brain-wide neurons. The analysis encompasses neuronal activity strength and neuronal correlation. The former includes the frequency, amplitude, and ratio of responsive neurons, while the latter includes the global correlation and intra-regional correlation. The latter analysis is further used to compare the degree of change before and after manipulating LC activity, as well as to assess the differences among various manipulations. (**B-D**) Intra- and inter-regional correlation changes induced by three control manipulations: ablation control (**B**), Opto-control (**C**), and Chemo-control (**D**). Left (i): MI matrices with change levels color-coded. The diagonal represents MI of intra-regional correlations, while the remaining ones represent MI of inter-regional correlations. Wilcoxon rank sum test was used to compare changes between before and after various manipulations, with corresponding *p* values indicated. Right (ii): averaged inter-regional correlations before and after various manipulations, represented by circle graphs. Only inter-regional connections with correlations exceeding the median value before manipulations are shown. (**Bi** and **Bii**) Data from the ablation control group. (**Ci** and **Cii**) Data from the Opto-control group. (**Di** and **Dii**) Data from the Chemo-control group.

**Figure S8.**
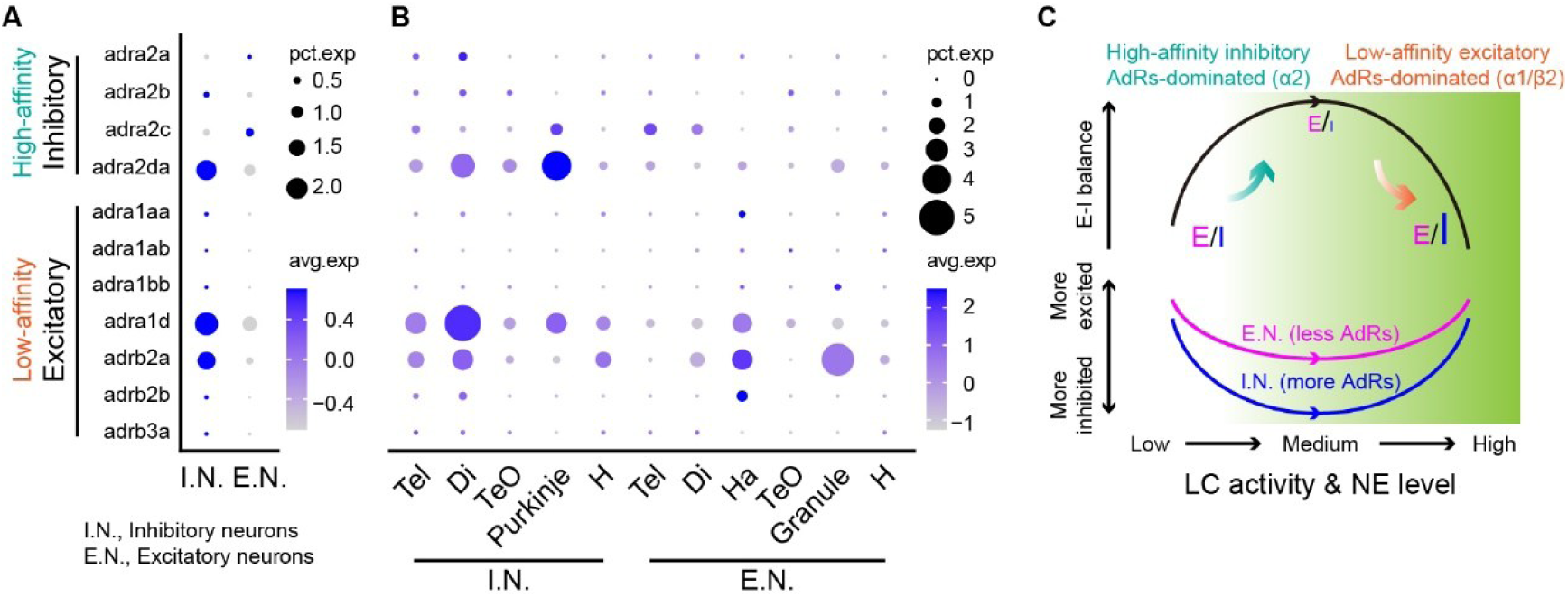
Brain-wide expression of NERs in inhibitory and excitatory neurons, related to Figures 7 and 8. (**A**) Dotplot showing the expression levels of excitatory and inhibitory noradrenergic receptors (NERs) in excitatory neurons (slc17a6a+/slc17a6b+, E.N.) and inhibitory neurons (gad1b+/gad2+/slc32a1+, I.N.). Percentage of cells expressing the gene (pct.exp) is size-coded, and average expression level (ave.exp) is color-coded. Transcriptomic data were from Zhang et al.^96^ (**B**) Dotplot showing the expression levels of excitatory and inhibitory NERs in inhibitory and excitatory across different brain regions, including the telencephalon (Tel), diencephalon (Di), optic tectum (TeO), hindbrain (H), habenula (Ha), and cerebellum (with Purkinje and granule cells representing inhibitory and excitatory neurons, respectively). Inhibitory neurons in the habenula are not shown due to their extremely low abundance.^98^ Percentage of cells expressing the gene (pct.exp) is size-coded, and average expression level (ave.exp) is color-coded. (**C**) Diagram of the potential mechanism underlying the LC inverted-U regulation of brain-wide neuronal activities by altering the excitatory/inhibitory (E/I) ratio at different NE levels.

## METHODS

### EXPERIMENTAL MODEL AND STUDY PARTICIPANT DETAILS

#### Zebrafish husbandry and preparations

Adult zebrafish (*Danio rerio*) were maintained in a circulating system at 28°C under a 14:10 hour light-dark cycle. All experiments were conducted using larvae at 5 - 7 days post-fertilization (dpf) at room temperature. All animal protocols were reviewed and approved by the Animal Care and Use Committee of the Center for Excellence in Brain Science and Intelligence Technology, Chinese Academy of Sciences (NA-046-2023).

#### Zebrafish lines

All transgenic zebrafish larvae used were in *nacre* background.^108^ The following published lines were used in this study: *Tg(UAS-E1b:Kaede)s1999t (abbreviation: Tg(UAS:Kaede))*,^109^ *Tg2(elavl3:GCaMP5G)ion13d (Tg(elavl3:GCaMP5G))*,^92^ *Et(gata2a:EGFP)pku2 (Et(vmat2:EGFP))*,^110^ *Tg(elavl3:NEXS-jRGECO1a)ion72d (Tg(elavl3:NES-jRGECO1a))*,^40^ *Tg(14×UAS-E1b:sypb-EGFP)ion7d (Tg(UAS:sypb-EGFP))*,^87^ *Tg(5×UAS-hsp70l)nkUAShspzGCaMP6s13a (Tg(UAS:GCaMP6s))*,^81^ *Tg(4×nrUAS:GFP)c369 (Tg(4×nrUAS:GFP))*,^111^ *Tg(UAS:mCherry)zf408 (Tg(UAS:mCherry))*,^112^ *Tg(elavl3:Hsa.H2B-GCaMP6f)jf7 (Tg(elavl3:H2B-GCaMP6f))*,^113^ *Tg(elavl3:GRAB_NE1m_)ion82d (Tg(elavl3:GRAB_NE1m_))*,^40^ *Tg(elavl3:GRAB_NE1h_)ion83d (Tg(elavl3:GRAB_NE1h_))*,^40^ *Ki(dbh:GAL4-VP16)ion36d (Ki(dbh:GAL4-VP16))*.^80^ In addition, *Tg(dbh:GAL4-VP16,5×UAS-hsp:ChrimsonR-tdTomato)ion120d (Tg(dbh:GAL4-VP16,UAS:ChrimsonR-tdTomato))* and *Tg(dbh:GAL4FF,5×UAS-hsp:Rno.TRPV1-TagRFP)ion88d (Tg(dbh:GAL4FF,UAS: TRPV1-TagRFP))* were constructed with a Tol2 system in this study.^114^

### METHOD DETAILS

#### Sparse labeling, imaging and reconstruction of individual LC-NE neurons

We crossed *Ki(dbh:GAL4-VP16);Tg(UAS:Kaede)*^80^ with *Tg(elavl3:GCaMP5G);Et(vmat2:EGFP)*, and screened for larvae with only one Kaede-labeled LC-NE neuron for subsequent morphological imaging and 3D reconstruction. GCaMP5G expressed in all neurons and EGFP expressed in monoaminergic neurons both were used as references for registration. Kaede exhibits green fluorescence prior to photoconversion and undergoes an irreversible transition from green to red fluorescence when exposed to ultraviolet light (approximately 350 - 400 nm). The larvae examined were reared from birth in an incubator that simulated natural light conditions (14:10 hour light-dark cycle). The ultraviolet components of the background illumination during rearing converted Kaede from green to red as early as 3 - 4 dpf, thereby enabling the visualization of LC-NE neuronal morphology.

*In vivo* imaging was performed in 6-dpf larvae using an upright NiE-A1 plus confocal microscope (Nikon, Japan) equipped with a 25×/1.1-NA water-immersion objective. The larvae were dorsally mounted in 1.5% low-melting point agarose (Invitrogen) dissolved in system water. GCaMP5G/EGFP and Kaede were sequentially excited using 488-nm and 561-nm lasers, respectively. Each 3D volumetric image was acquired with a resolution of 1024 × 1024 pixels (0.488 μm/pixel), covering a depth of 290 μm at 1-μm z-step. To image the complete axon projections of LC-NE neurons, three overlapping volumes (5% overlap) from the telencephalon to the spinal cord were acquired and automatically stitched. In addition, high-resolution imaging (4× zoom) was performed to resolve the dense arborization around the LC-NE soma. The morphology was reconstructed semi-automatically using the Simple Neurite Tracer plugin in ImageJ (NIH), with each reconstruction independently verified by at least two researchers.

#### Discrimination of LC-NE neuronal dendrites and axons

To discriminate dendrites and axons of LC-NE neurons, we used *Ki(dbh:GAL4-VP16);Tg(UAS:mCherry);Tg(UAS:sypb-EGFP)* larvae, in which the varicosities along the LC-NE neuronal axon are labeled with synaptophysin b (sypb), a synaptic vesicle protein, fused to EGFP, and the overall morphology of LC-NE neurons are labeled with mCherry. As sypb-EGFP localizes specifically to presynaptic axon terminals, LC-NE neuronal dendrites were identified by lack of EGFP-positive puncta. Based on this strategy, we observed that LC-NE neuronal dendrites are relatively short and proximal to the soma, and terminate within the ventrolateral perisomatic region. We analyzed four representative LC-NE neurons, all of which exhibited consistent features. Leveraging these morphological criteria, we then manually classified the dendrites and axons of all 165 reconstructed LC-NE neurons.

#### Clustering analysis of the morphology of individual LC-NE neurons

To classify the morphological types of LC-NE neurons, all right hemisphere LC-NE neurons were mirrored to the left hemisphere along the midline of the brain template. A total of 165 reconstructed LC-NE neurons were subsequently subjected to morphological clustering using NBLAST, a rapid and sensitive algorithm for quantifying pairwise neuronal similarity.^88^ The clustering workflow was implemented in R using the nat and nat.nblast packages. Firstly, the morphology of individual LC-NE neurons was imported using the read.neurons function. Secondly, each neuron’s morphology was converted into a directed point cloud using the dotprops function with parameters resample = 1 and k = 2, generating a lightweight geometric abstraction of its morphology. Thirdly, pairwise similarity scores were computed using the nblast_allbyall function, which performs all-by-all comparisons of neuronal morphology (https://github.com/jefferislab/nat.nblast). Fourthly, the resulting similarity matrix was subjected to hierarchical clustering using the nhclust function (https://github.com/jefferis/nat). Fifthly, clustered neuronal morphologies were visualized using the plot3d function. Finally, the clustering results were manually checked by two independent researchers to form the final clusters.

#### Brain template and registration

To establish a standardized spatial framework for quantifying individual LC-NE neurons’ morphology, we developed a brain template using 6-dpf *Tg(elavl3:GCaMP5G);Et(vmat2:EGFP)* larvae. From eleven imaged seed brains, we selected three candidates as reference brains. Template generation involved nonrigid registration of the remaining brains to each reference using the Computational Morphometry Toolkit (CMTK-3.2.3; http://www.nitrc.org/projects/cmtk/), implemented with the following command string: -awr 010203 -∣fa -g -T 24 -X 52 - C 8 -G 80 -R 3 -E 1 -A ‘--accuracy 0.4’ -W ‘--accuracy 1.6’.^115^ The three candidate references were quantitatively evaluated by their normalized cross-correlation (NCC), mean square distance (MSD), mean absolute distance (MAD), and cross-entropy (Hxy). The reference brain demonstrating optimal alignment metrics (highest NCC and lowest MSD, MAD, and Hxy) was selected for template construction. This optimal reference was then intensity-averaged with all aligned seed brains to generate the brain template.^115^ The brain of individual larvae, each labeled with a single LC-NE neuron, was then registered to the brain template based on their corresponding GCaMP5G/EGFP signals using Advanced Normalization Tools (ANTs).^116^

To leverage the existing brain region annotation information from the Zebrafish Brain Mesoscale Atlas,^86^ which used 6-dpf *Tg(elavl3:H2B-GCaMP6f)* larvae to generate the brain template, we established a spatial correspondence between our *Tg(elavl3:GCaMP5G);Et(vmat2:EGFP)* template and the annotated *Tg(elavl3:H2B-GCaMP6f)* template. We generated a brain template for *Tg(elavl3:NES-jRGECO1a);Tg(elavl3:H2B-GCaMP6f)* larvae to serve as a bridge template. As the pan-neuronal and cytoplasmic expression patterns of *Tg(elavl3:GCaMP5G)* and *Tg(elavl3:NES-jRGECO1a)* are similar, the bridge template enables accurate transformation between templates. The transformation matrices were subsequently applied to map reconstructed individual LC-NE neurons’ morphologies onto the annotated template, enabling brain region-specific analysis.

#### *In vivo* whole-cell recording and sensory stimulation

To characterize the physiological properties of LC-NE neurons, we performed *in vivo* whole-cell recording following morphological imaging (see Figure S2A). Larvae were paralyzed by 5-min immersion in the neuromuscular junction blocker α-Bungarotoxin (100 μg/ml, Tocris Bioscience) and then embedded in 1.5% low-melting point agarose. LC-NE neurons, located within the rhombomeres 1 and 2 (R1 and R2) of the hindbrain, were clearly visualized under fluorescence and bright field illumination. The gel covering the hindbrain was removed, and the skin overlying the ventricle was dissected to facilitate electrode penetration into the brain tissue. Perfusion with extracellular solution (in mM: 134 NaCl, 2.9 KCl, 2.1 CaCl_2_•2H_2_O, 1.2 MgCl_2_•6H_2_O, 10 HEPES, 10 glucose; 290 mOsm, pH = 7.8) maintained physiological conditions. Recording micropipettes (20 - 30 MΩ, 1 - 1.5 μm tip diameter) pulled from borosilicate glass capillaries (BF100-58-10, Sutter Instrument, USA) were filled with an intracellular solution (in mM: 100 K-gluconate, 10 KCl, 2 CaCl_2_•2H_2_O, 2 Mg_2_•ATP, 0.3 GTP•Na_4_, 2 phosphocreatine, 10 HEPES, 10 EGTA; 270 mOsm, pH = 7.4). LC-NE neurons were recorded with an EPC-10 amplifier (HEKA, Germany). After the formation of the giga seal, a short negative pressure was applied to break the cell membrane beneath the micropipette tip. The equilibrium potential for Cl^−^ was about -60 mV, as calculated using Nernst equation. The signals were filtered at 2.9 kHz and sampled at 10 kHz.

Membrane capacitance was measured during membrane capacitance compensation using the electrophysiological recording software. In voltage-clamp mode, the cell was held at -60 mV. To examine the current-voltage (I-V) curve, 1-s voltage steps were applied, ranging from -100 to 40 mV with a 10-mV step. Membrane resistance was calculated as the reciprocal of the slope of the linear fit to the whole-cell current recorded in voltage-clamp mode with the holding potential from -100 to -60 mV. Subsequently, the recording was switched to current-clamp mode, current steps ranging from -40 to 100 pA were injected into the recorded neuron at a 10-pA step, each lasting 1 s.

To characterize the salience-response properties of LC-NE neurons, we employed four distinct sensory stimuli. For sensory responses recorded in current-clamp mode (I = 0 pA), we applied three types of sensory stimuli: (1) a 2-s full-field white light flash was produced by a LED controlled by a Master-8 pulse generator (A.M.P.I, Israel); (2) looming stimuli, which mimic the expanding shadows of approaching predators, were presented using a projector;^82^ (3) auditory responses were evoked by a 10-ms pure tone at 500 Hz and 85 dB.^83^ In addition, mechanical stimulation was incorporated for calcium imaging-based salience assessment (see Figures 5I and 5K). For mechanical stimulation, the extracellular solution was locally puffed at the trunk region (proximal to the 6^th^ myotome with local agarose removed) using a glass micropipette (tip diameter: ∼10 μm) positioned ∼10 μm above the skin surface. Puff stimulation was controlled by a Master-8 pulse generator (A.M.P.I., Israel) interfaced with a pressure ejection system (Picospritzer III, Parker, USA). Puff (12 psi, 200-ms duration) was repeated four times at a 30-s interval. The response was quantified by measuring action potential (AP) numbers and firing rates, both of which were averaged across 4 - 10 trials within stimulus-specific analysis time windows.

In dual whole-cell recordings, spontaneous activities of the paired LC-NE neurons were recorded in current-clamp mode (I = 0 pA) in 6-dpf *Ki(dbh:GAL4-VP16);Tg(4×nrUAS:GFP)* larvae. To test electrical coupling, 0.5-s current pulses (± 50 pA, ±100 pA) were injected into one neuron while recording both cells’ responses.

#### Clustering analysis of physiological properties of LC-NE neurons

To examine whether morphologically defined LC-NE neurons exhibit distinct functional characteristics, we performed a cluster analysis using k-means clustering on various parameters related to their intrinsic electrophysiological properties and sensory-evoked responses, followed by the gap statistic analysis to determine the optimal number of clusters.^89^ The within-cluster dispersion of the observed data was compared to that of a reference null distribution. This null distribution was generated by repeatedly sampling (100 times) from a uniform distribution across the range of our experimental measurements. The expected dispersion was defined as the average log of within-cluster dispersion across these replicates. For each potential cluster number *k*, we calculated the gap statistic Gap(*k*) as the difference between the expected and observed log of dispersion. Following the original criterion, the optimal number of clusters *k* was selected as the smallest *k* such that Gap(*k*) ≥ Gap(*k*+1) − *s_k_*_+1_. That is, the increase in the gap statistic from *k* to *k*+1 did not exceed the “1-standard-error” of the gap statistic at *k*+1. The outcome of *k* = 1 indicates no significant functional characteristic segregation among LC-NE neurons, whereas *k* > 1 would reveal functionally distinct groups.

We applied the aforementioned procedure to analyze 28 examined LC-NE neurons, which have defined morphological classifications and electrophysiological data. Each neuron was characterized by 11 electrophysiological features: resting membrane potential, membrane capacitance, input resistance, Boltzmann sigmoidal slope of the current-voltage (I-V) relationship, the number of APs evoked by a 100-pA current injection, as well as the AP number and frequency of responses to flash, looming, and pure tone stimuli. The results showed that the first *k* satisfying Gap(*k*) ≥ Gap(*k*+1) - *s_k_*_+1_ was 1, indicating an optimal cluster number of 1 (see Figure S2F). Although the gap values exhibited a slight increasing trend with larger *k*, the increments consistently remained below one standard error (*s_k_*_+1_). Moreover, all values of Gap(*k*+1) − *s_k_*_+1_ either fell within or below the confidence interval defined by Gap(*k*) − *s_k_*, suggesting a statistically insignificant clustering tendency. We therefore selected *k* = 1 as the optimal number of clusters, indicating no significant functional segregation among LC-NE neurons. Then, we performed dimensionality reduction using both principal component analysis (PCA) and t-distributed stochastic neighbor embedding (t-SNE) on the 11-dimensional parameter space. Both methods consistently revealed diffuse distributions across 28 LC-NE neurons examined, further confirming their comparable electrophysiological and sensory response properties.

#### Single-cell SMART sequencing and analysis of individual LC-NE neurons

LC-NE neurons were isolated from 6-dpf *Ki(dbh:GAL4-VP16);Tg(4×nrUAS:GFP)* larvae for single-cell transcriptomics.^117^ To specifically target LC-NE neurons in R1 and R2 while excluding medulla oblongata noradrenergic (MO-NE) neurons in R8, we microdissected the hindbrain tissues containing LC-NE neurons (see Figure 3A) using ophthalmic scissors under a stereomicroscope, following anesthesia of the larvae with MS-222. Tissue was enzymatically dissociated in 200 µL papain solution at 37°C for 15 min, quenched with 800 µL wash buffer, and subsequently concentrated to 50 - 100 µL by centrifugation. Single-cell suspension was blown via tubes of different calibers and then spread onto a gel plate. Individual GFP-positive neurons were manually selected using a glass electrode (inner diameter: 30 µm) and transferred to individual PCR tubes containing RNA-free buffer solution for SMART sequencing. All sample preparations were completed within 1.5 hours. Subsequent steps followed the SMART-seq2 protocol (Takara Bio). Indexed libraries were sequenced on Illumina NovaSeq 6000 using a paired-end 150 bp (PE150) strategy. Across four experiments involving approximately 130 larvae, we collected data from 215 LC-NE neurons. Sequencing achieved a depth of over 1 million uniquely mapped reads per cell, with a median of 13.3 million reads. Samples with fewer than 1 million uniquely mapped reads (6 cells) were excluded from further analysis.

Paired-end reads were aligned to the *Danio rerio* genome (GRCz11, v.99; Ensembl) using HISAT2 (v.2.1.0), achieving an average mapping rate of 87.1% (Table S3). The mapping output in SAM files were directed to BAM files using SAMtools (v.1.6). Transcript expression was quantified using Subread (v.2.0.1) based on GTF annotations and BAM alignments, resulting in 209 LC-NE neurons for subsequent analysis. Single-cell raw count matrices for these neurons were imported into R (v.4.2) and processed using the Seurat package (v.4.4.0). Low-quality and dying cells were filtered out based on unique feature counts (< 200 or > 2,500) and mitochondrial content (>5%). This filtering resulted in a final dataset of 209 neurons with a median of 7,200 detected genes per cell and a median mitochondrial content of 0.9%. The filtered data were log-normalized and scaled using default parameters. PCA was performed on the scaled data, and significant principal components (PCs 1 - 4, 7, 10, 35; *p* < 0.0001) were used for graph-based clustering. Non-linear dimensionality reduction via t-SNE visualized three distinct LC-NE neuron clusters at a resolution of 0.6. Different resolution conditions were tested using R package ROGUE (v.1.0). Following that, we used ‘FindAllMarkers’ algorithm to find significant differential expression of marker genes (Table S4) (*p*_val_adj < 0.0001). Gene Ontology (GO) analysis of marker genes for the three LC-NE neuron clusters was conducted using the clusterProfiler package (v.4.10.1). Biological process (BP) terms were used for analysis.

To functionally characterize the three LC-NE neuron clusters, we collected data on 654 genes (Table S2) related to LC-NE neuronal function and development based on literature and ZFIN database (https://zfin.org/). These genes are associated with sodium channels (n = 15), potassium channels (n = 84), calcium channels (n = 44), chloride channels (n = 11), gap junctions (n = 44), neurotransmitter biosynthesis (n = 17), neurotransmitter release (n = 35), neurotransmitter transporters (n = 47), neurotransmitter receptors (n = 144), neuropeptides (n = 45), neuropeptide receptors (n = 55), developmental markers (n = 9), cytoskeletal organization (n = 37), and axon extension and guidance (n = 67). Z-score heatmaps of 202 genes expressed in >30% of LC-NE neurons across the three clusters (Table S5) were generated using the ComplexHeatmap package (v.2.18.0). To identify LC-NE neuron clusters with active gene sets, we used the “Area Under the Curve” (AUC) to calculate whether a critical subset of the input gene set is enriched among the expressed genes for each cluster using AUCell package (v.1.24.0). The distribution of AUC scores across clusters was compared using one-way ANOVA.

#### Immunohistochemistry

Whole-mount immunostaining of tyrosine hydroxylase (Th) was performed in 6-dpf *Ki(dbh:GAL4-VP16);Tg(4×nrUAS:GFP)* larvae. Larvae were fixed with 4% paraformaldehyde (PFA) in phosphate-buffered saline (PBS) at 4°C overnight. Brains were dissected using fine forceps and subsequently washed three times with PBST (PBS containing 1% Tween-20) for 5 min per wash. Permeabilization was achieved using the following sequential steps: 5 min in 75% PBST + 25% methanol at room temperature (RT), 5 min in 50% PBST + 50% methanol at RT, 5 min in 25% PBST + 75% methanol at RT, 4 hours in pre-cooling 100% methanol at -20°C, 5 min in 25% PBST + 75% methanol at RT, 5 min in 50% PBST + 50% methanol at RT, 5 min in 75% PBST + 25% methanol at RT, 5 min in PBST for 3 times at RT, 5 min in ddH_2_O at RT, 10 min in pre-cooling 100% acetone at -20°C, 5 min in PBST for 3 times at RT. After the permeabilization, samples were blocked with commercial blocking solution (QuickBlock™ Blocking Buffer for Immunol Staining, Beyotime, P0260) at 4°C overnight. Primary labeling was done by incubating samples into the commercial blocking solution with anti-Th (1:200, Anti-tyrosine hydroxylase, mice, Millipore, MAB318) at 4°C for 48 hours. Samples were then washed twice with PBST for 1 hour each, followed by two additional 15-min washes. Secondary labeling was done by incubating samples into commercial blocking solution with Alexa Fluor^TM^ 568 goat anti-mouse secondary antibodies (1:500, Molecular Probes) at 4°C for 36 hours. Samples were then washed three times with PBST for 10 min each and once for 30 min. Imaging was conducted using an FV3000 confocal microscope (Olympus, Japan).

#### Neurobiotin-based gap junction tracing

To examine gap junctions among LC-NE neurons, neurobiotin (2.5%) was loaded into the recorded LC-NE neuron of 6-dpf *Ki(dbh:GAL4-VP16)*;*Tg(4×nrUAS:GFP)* larvae via a whole-cell recording micropipette for over 30 min. Following loading, the larva was allowed to recover for 2 - 3 hours to permit neurobiotin diffusion through gap junctions. The larva was then fixed overnight at 4°C in 4% PFA. After optimized permeabilization and immunostaining,^92^ the sample was embedded in 1.5% low-melting point agarose for imaging. Imaging was performed using an FV3000 confocal microscope (Olympus, Japan) with excitation of 488 nm for GFP and 559 nm for streptavidin-conjugated Alexa Fluor^TM^ 594.

#### *In vivo* calcium imaging and analysis of LC-NE neurons’ and brain-wide neurons’ activities

To monitor population LC-NE neurons’ activities, we performed *in vivo* confocal calcium imaging on 6-dpf *Ki(dbh:GAL4-VP16);Tg(UAS:GCaMP6s)* larvae using an upright FV1000 confocal microscope (Olympus, Japan) equipped with a 40×/0.8-NA water-immersion objective. Spontaneous activities of bilateral LC-NE neuronal somata were recorded at approximately 6 Hz for 10 min in paralyzed larvae. Image sequences were motion-corrected using ImageJ (NIH), followed by the manual selection of somatic regions of interest (ROIs) to extract original gray values (F). Fluorescence traces were analyzed in MATLAB (MathWorks) by calculating ΔF/F_0_ = (F−F_0_)/F_0_, where F_0_ represents the baseline fluorescence intensity.

To investigate how LC-NE neurons influence downstream neural networks, we acquired 10-min brain-wide spontaneous calcium activities before and after manipulations of LC-NE neurons (see Figure 7A). For LC ablation and LC chemogenetic activation experiments, an Ultima Investigator two-photon microscope (Bruker, USA) equipped with an Olympus 20×/0.95-NA water-immersion objective was used. The laser was tuned to 920 nm, and imaging was conducted at a frame rate of 2.3 Hz, with a resolution of 0.67 μm/pixel in resonant mode. For LC optogenetic activation experiments, an Opterra confocal microscope (Bruker, USA) equipped with a Nikon 16×/0.8-NA water-immersion objective was used. The laser was tuned to 488 nm, and imaging was conducted at a frame rate of 2.5 Hz, with a resolution of 0.49 μm/pixel. Two imaging layers were acquired during brain-wide calcium imaging: one layer captured LC-NE neurons, and the other encompassed regions spanning from the telencephalon to the hindbrain (see Figure 7B), yielding 2,000 - 3,000 neurons per larva. Larvae were immobilized with α-Bungarotoxin to eliminate motion artifacts during imaging.

The analysis of brain-wide calcium activities was carried out following the workflow outlined in Figure S7A. First, single-neuronal ROIs were automatically extracted from the averaged time-course imaging with custom codes. After removing the background noise, calcium signals of each neuron were detected along the time course and used to calculate ΔF/F_0_. We analyzed the frequency and amplitude of each neuron’s activities, and ratio of responsive neurons in whole time window, and then analyzed intra- and inter-regional neuronal correlation with custom codes (Figure S7A). The first 100 frames (approximately 43 s) of data from two-photon imaging were excluded from analysis to mitigate potential artifacts introduced by the initial up-and-down movement of the piezo. For confocal imaging, the first 80 - 160 s of data were excluded to eliminate the effects of the startup of the 488-nm laser, which could evoke light responses of LC-NE neurons, and initial piezo movement. Consistent data treatment procedures were applied to the control groups.

To quantitatively compare LC-NE neuron activities under different manipulations, ROIs for LC-NE neurons were manually identified from aforementioned brain-wide calcium imaging data to extract original gray values, followed by ΔF/F_0_ calculation and detection of significant responses by custom codes. ΔF/F_0_ was transformed into Z-score in order to perform normalization and compare different functional profiles derived from one- or two-photon microscope. The baselines of Z-score curves were corrected to avoid negative values. The area under the baseline-corrected Z-score curves (Z-score AUC) within specific time windows was calculated. For optogenetic data, time window began 40 s after the startup of the 561-nm laser to eliminate the contribution of visual responses to neuronal activities (see Figures 6C and 6D). For chemogenetic data, time window commenced with the onset of LC-NE neurons’ responses to capsaicin. Only Z-score AUC of significant responses contributed to final results for statistical analysis.

#### *In vivo* imaging of NE release

To observe brain-wide NE release, we performed *in vivo* time-lapse confocal or light-sheet imaging in paralyzed *Tg(elavl3:GRAB_NE1h_)* larvae at 6 dpf. To observe local NE release within the TeO_NL at a fine scale, we performed *in vivo* time-lapse confocal imaging in paralyzed *Tg(elavl3:GRAB_NE1m_)* larvae at 6 dpf. These GRAB sensors enabled sensitive detection of endogenous NE release.^40^ For brain-wide spontaneous NE release, a customized light-sheet microscope was used with 488 nm excitation at a volumetric acquisition rate of 2 Hz. Imaging was performed across a volume of 25 layers with an 8-μm step interval and lasted for 3 - 4 min. For looming-evoked NE release, confocal imaging was conducted using a FN1 microscope (Nikon, Japan) equipped with a 16×/0.8-NA water-immersion objective. Imaging was performed at a 10-μm step interval across a depth of approximately 120 μm, achieving a volumetric imaging rate of 0.5 Hz. Looming stimuli with similar parameters described in Figure S2A were presented in front of the larva via a projector. Procedures similar to those used for calcium activity processing, as mentioned above, were applied to analyze NE release signals. We performed *in vivo* imaging of NE release in the optic tectum following optogenetic and chemogenetic manipulations of LC-NE neurons to assess NE release levels (Figures S6G-S6I). The data were processed in the same manner as the LC-NE calcium imaging data.

#### Two-photon laser-based ablation of LC-NE neurons

To assess LC effects on brain-wide neuronal activities, two-photon ablation of bilateral LC-NE neurons was performed in 6-dpf *Tg(dbh:GAL4FF,UAS:TRPV1-TagRFP)*;*Tg(elavl3:H2B-GCaMP6f)* larvae, using an Ultima Investigator two-photon microscope (Bruker, USA) equipped with a 20×/0.95-NA water-immersion objective. We used 800-nm laser to precisely ablate LC-NE neurons which were marked by RFP. Successful ablation was confirmed by both the appearance of bulb-like structures under brightfield microscopy and complete loss of RFP signal. Control ablations targeted an equivalent number of randomly selected GCaMP6f-expressing neurons around the LC soma region.

To distinguish projection patterns of LC-NE neurons and MO-NE neurons, 800-nm laser-based ablation of bilateral LC-NE neurons was conducted in *Ki(dbh:GAL4-VP16);(Tg(4×nrUAS:GFP)* larvae at 5 dpf, using an upright FVMPE-RS multi-photon microscope (Olympus, Japan), and fluorescent imaging was performed at 6 dpf to examine the remaining projection pattern of MO-NE neurons.

#### Optogenetic activation of LC-NE neurons

Optogenetic activation of bilateral LC-NE neurons was performed in 6-dpf *Tg(dbh:GAL4-VP16,UAS:ChrimsonR-tdTomato)*;*Tg(elavl3:H2B-GCaMP6f)* larvae using an Opterra confocal microscope (Bruker, USA). We designed a seamless protocol combining pre-optogenetics imaging, LC optogenetics, and post-optogenetics imaging to assess brain-wide neuronal activities. Initially, a 10-min recording of spontaneous brain-wide neuronal activities was performed using a 488 nm laser at 2.5 Hz. Subsequently, based on pre-defined parameters, the ROIs for optogenetic stimulation were precisely targeted as a 30-µm diameter circle centered on the soma region of bilateral LC. The LC soma region was segmented into three layers from dorsal to ventral with a 10-µm step interval for sequential activation. LC-NE neurons were optogenetically activated using 561-nm laser stimulation (30% power) targeting ChrimsonR. Stimulation consisted of 200-ms pulses delivered at 2.5 Hz, repeated ten times per layer, resulting in a total stimulation duration of 12 s for all LC-NE neurons. Immediately following stimulation, a second 10-min recording of spontaneous brain-wide neuronal activities was conducted to capture post-optogenetics neuronal activities. LC-NE neuronal activities were continuously monitored to verify the efficiency of the optogenetic activation protocol. For control experiments, we used larvae in which LC-NE neurons expressed mCherry but not ChrimsonR. The same experimental procedures were conducted.

#### Chemogenetic activation of LC-NE neurons

Chemogenetic activation of LC-NE neurons was performed in 6-dpf *Tg(dbh:GAL4FF,UAS:TRPV1-TagRFP)*;*Tg(elavl3:H2B-GCaMP6f)* larvae, using an Ultima Investigator two-photon microscope (Bruker, USA). TRPV1 channels were specifically expressed in NE neurons and activated by application of capsaicin. Prior to the chemogenetic activation of LC-NE neurons, we imaged spontaneous brain-wide neuronal activities for 10 min using a 920 nm laser. Capsaicin, dissolved in DMSO, was then bath-applied to the sample dish, achieving a final concentration of 5 µM. LC-NE neurons typically responded on average within 1 min after capsaicin application. Imaging resumed immediately after capsaicin application, capturing spontaneous activities for an additional 10 min at the same imaging planes. Concurrently, LC-NE neuronal activities were continuously monitored to verify the activation onset and efficiency during the chemogenetic protocol. For control experiments, DMSO was applied alone, without capsaicin. Please note, the bath application of capsaicin may supposedly activate MO-NE neurons, because these cells also express TRPV1. The two-photon ablation of bilateral LC-NE neurons revealed that MO-NE neuronal projections are confined primarily to the ventral diencephalon, ventral hindbrain, medulla, and spinal cord, and are spatially segregated from LC-NE neuronal projections (see Video S4). Notably, none of the regions of interest for LC-related NE release and functional analyses overlap with projection territories of MO-NE neurons.

#### Calculation of the turning point of the inverted-U relationship

To characterize the inverted-U relationship between the level of LC-NE neuronal activities and the changes of brain-wide neuronal activities, the data in Figure 8D were used to estimate the turning point of the inverted-U curve, which represented the modulation index of intra-regional correlations within specific brain regions *versus* log2-transformed level of LC-NE neuronal activities corresponding different experimental manipulations. As the logarithm base 2 of zero approaches negative infinity, the data of the LC ablation group, in which there was absolutely no activity of LC-NE neurons, were excluded. As the tendency of the inverted-U curve is asymmetrical and exhibit nonlinear changes, the remaining data were fitted to a third-order polynomial regression: y = *p*_1_ × x^3^ + *p*_2_ × x^2^ + *p*_3_ × x + *p*_4_. Ridge regression with λ = 0.001 was applied to avoid overfitting. The x-value, at which the first derivative equals zero and the y-value is maximized, indicates the turning point. These turning points were subsequently correlated with the density index of population LC-NE neurons’ projections in each brain region. The fitting results were listed below:

|  |  |
| --- | --- |
| Tel | $y = 0.0069 \times x^3 - 0.1352 \times x^2 + 0.7927 \times x - 1.3056$ |
| | Turning point = 4.4570, $R^2 = 0.9986$, RMSE = 0.0032 |
| Ha | $y = -0.0108 \times x^3 + 0.1470 \times x^2 - 0.5715 \times x + 0.6699$ |
| | Turning point = 6.2909, $R^2 = 0.9996$, RMSE = 0.0024 |
| TeO_PL | $y = -0.0134 \times x^3 + 0.1804 \times x^2 - 0.6551 \times x + 0.6965$ |
| | Turning point = 6.4208, $R^2 = 0.9997$, RMSE = 0.0027 |
| CCe | $y = -0.0126 \times x^3 + 0.1788 \times x^2 - 0.7243 \times x + 0.8773$ |
| | Turning point = 6.5004, $R^2 = 0.9994$, RMSE = 0.0030 |
| H | $y = -0.0020 \times x^3 + 0.0019 \times x^2 + 0.1658 \times x - 0.4422$ |
| | Turning point = 5.5466, $R^2 = 1.0000$, RMSE = 0.0007 |

### QUANTIFICATION AND STATISTICAL ANALYSIS

One-way multivariate analysis of variance (one-way MANOVA) was performed for pairwise comparisons between clusters (Figure 1E).

For multiple-group comparisons for normally distributed data, one-way ANOVA was performed (Figures 2C-2E, 2G, 2I-2L; Figures S2C-S2E; Figure 3D; Figure S3G; Figure 4I; Figure S5H). For non-normally distributed data, Friedman test (Figures 6C and 6J; Figure 7G) or Kruskal-Wallis test (Figure 8D) was performed for matched and independent samples, respectively.

For two-group comparisons for normally distributed data, unpaired Wilcoxon rank sum test (Figures S1G-S1I), unpaired t-test (Figures 1F-1I; Figures S1P and S1Q; Figures 4B, 4D and 4G; Figures S4B and S4D; Figure 5C) or paired t-test (Figures S1P; Figures 4C and 4F; Figures S4A and S4C; Figures S5B, S5C and S5G; Figures 6F, 6H and 6O) was performed for independent and matched samples, respectively.

For two-group comparisons for non-normally distributed data, Mann-Whitney test (Figures S6F and S6I; Figures 7Hi, 7Hiii; Figures S7Bii, S7Dii; Figures 8A-8C) or Wilcoxon matched-pairs signed-rank test (Figures 4C, 4F, 4I; Figures S4A, S4C; Figures 6C, 6J, 6O; Figure 7Hii; Figure S7Cii) was performed for independent and matched samples, respectively.

For comparisons of the distributions between two groups, two-sample Kolmogorov-Smirnov test (Figures S1J-S1L; Figures 2F and 2H) was performed.

For comparisons with zero, one-sample t-test (Figures 7C-7F; Figures 8A-8C) or one-sample Wilcoxon signed rank test (Figures 7C-7F; Figures 8B, 8C) was performed for normally and non-normally data, respectively.

For ordinary least squares linear regression, the dotted lines represent the 95% confidence bands of the best-fit line. A t-test was conducted to evaluate whether the slope significantly deviates from zero (Figure 6D; Figure 8E).

Normality was assessed using Kolmogorov-Smirnov test or D’Agostino & Pearson omnibus test.

n.s., not significant; **\****p* < 0.05, **\*\****p* < 0.01, **\*\*\****p* < 0.001, **\*\*\*\****p* < 0.0001. Data are represented as mean ± SEM.

## Notes

### Competing Interest Statement

The authors have declared no competing interest.

